# Fermentation of Agricultural By-products and Zinc Supplementation: A Synergistic Approach to Mealworm Microbiome Optimization

**DOI:** 10.1101/2025.08.05.668463

**Authors:** B. Antunes, C. Zanchi, M. Stoffregen, S. Grebenteuch, M. Bogdanova, S. Mbedi, S. Sparmann, H. Haase, J. Rolff, C. Rafaluk, C. Keil

## Abstract

The increasing world population has put pressure on food industry to swiftly come up with initiatives that can sustain an increase in the demand for food, which have led to environmentally unsustainable practices. Insects, such as *Tenebrio molitor* (TM) larvae, have been widely proposed as sustainable alternative sources of protein with less environmental footprint. Insects can be reared with diets featuring low-cost agricultural side-streams, promoting circular economical practices, which can also be fermented to promote the development of probiotic microbiota. The inclusion of probiotics in the diets of insects during rearing has recently been explored to mitigate potential exposure to pathogens, potentially serving as alternatives to antibiotics. The focus of this study was the fermentation of spinach to naturally develop complex probiotic populations to enrich the diet of mealworms, promoting host health, thus resulting in a more nutritious food/feed source, featuring a safer microbiome profile.

Sequencing revealed that fermented spinach had increased lactic acid bacteria (LAB), while repressing *Pseudomonas*. LAB species have been acknowledged and used as probiotics, whereas *Pseudomonas* populations are often associated with opportunistic resistant infections. Moreover, zinc is an essential element for bacterial microbiomes of many organisms, including insects. As the effects of dietary zinc on insect gut microbiome have been scarcely investigated, we have supplemented our formulated diets with this trace element and fed it to TM larvae for 35 days. While various diet compositions did not affect mealworm mortality, larvae fed spinach-based and zinc-rich diets gained less weight than that of control. Furthermore, higher dosage of zinc-supplementation resulted in significantly higher accumulation in TM larvae and a lower bioaccumulation factor. Sequencing of larvae fed diets supplemented with fermented spinach revealed increased presence of probiotic-associated genera, *Lactococcus* and *Weissella*. It was also found that supplementing diets with spinach and zinc significantly modulated the microbiome of larvae.

## Introduction

The increasing world population and corresponding increase in the demand for food has put pressure on food production, promoting unsustainable harvesting practices across the planet, leading to habitat loss, deforestation, animal overexploitation and increasing greenhouse gas emissions (Ordoñez-Araque and Egas-Montenegro, 2021). Consequently, innovative solutions, food alternatives and sustainable development of agricultural and food systems are required. The Food and Agriculture Organization (FAO) of the United Nations, has proposed insects as an environmentally friendly alternative food and feed source, to ensure food security for the growing world population (van Huis, 2021). Insect rearing produces less greenhouse gases and ammonia emissions, requires less water than conventional protein sources and can be produced in vertical farming systems, leading to a lower space requirement (Noyens et al., 2023). Furthermore, it has been shown that insects are able to convert low-value biowaste into valuable protein-rich biomass (Oonincx et al., 2015). The European Union has established the utilization of insects for food (EU regulation 2015/2283) and feed (EU 2017/893) (EU Commission, 2015; 2017).

Amongst the insects evaluated, the yellow mealworm (*Tenebrio molitor* larvae, Coleoptera: Tenebrionidae) has received extensive attention because of its high macronutrient value (Rumpold and Schluter, 2013) and its wide range of potential food and feed applications (Gkinali et al., 2022; Khanal et al., 2023). Because of its low mobility, larvae and adults can be easily reared (Morales-Ramos et al., 2024; Ortiz et al., 2016). Moreover, mealworms can be grown using low grade organic bioresources or byproducts, leading to higher feed conversion efficiency, while requiring less water and land for growth, potentially minimizing greenhouse gases (Adamaki-Sotiraki et al., 2024; Grau et al., 2017; Kotsou et al., 2024). *T. molitor* (TM) is also well known as a model organism for studies of innate immunity (Evison et al., 2017; Sadd et al., 2006), and a full genome assembly has been published (Mann et al., 2024; Oppert et al., 2023).

When used as food and feed, TM larvae are mildly processed and eaten mostly without removal of their gut or any other part of their body, thus are considered as natural carriers of microorganisms (Osimani et al., 2018). One strategy that has been explored to mitigate potential exposure to pathogens, parasites and toxins from insect consumption, is the integration of probiotics in the diets of insects during rearing (Grau et al., 2017; Savio et al., 2022). The idea of probiotic prophylaxis emphasizes the usage of competitive elimination for enhancing a particular ecosystem (Krishna et al., 2022). The FAO and WHO describe probiotics as “live micro-organisms which when administered in adequate amounts confer a health benefit on the host” and they are considered as environmentally friendly biocontrol agents for usage in agriculture and food industry (Hossain et al., 2017; Hotel and Cordoba, 2001). Administration of probiotics has been shown to improve health, growth rate, productivity and even disease resistance of animals, serving as alternatives to antibiotic use (Arsene et al., 2021). Although increasing in popularity, the data on the effects of the administration of probiotics to TM larvae are still limited and focus on the addition of probiotic cultures to the diet (Häbermann *et al*., 2025; Lecocq *et al*., 2021; Rizou *et al*., 2022; Savio *et al*., 2024; Zhong *et al*., 2017).

Gut microbiota plays a key role in the health, productivity and well-being of the insect host, influencing nutrient absorption, metabolism, and immune function. Thus, optimizing insect’s diets to provide nutrients both to the insects and their microbiota could produce insects that are nutritiously more balanced and safer for the final consumer (Liberti and Engel, 2020). Notably, bacteria in the microbiome produce beneficial metabolites (e.g., short-chain fatty acids), which reduce inflammatory markers and lower intestinal luminal pH, thus limiting the proliferation of potentially pathogenic bacteria (Macfarlane and Macfarlane, 2012; Martin-Gallausiaux et al., 2021). A common method of enhancing the nutritional value of insects is through their diet, featuring high-density of the desired component, called gut-loading, supplementing the nutrients contained in the insect’s body (Finke, 2003).

Although TM larvae are rich in vitamins and minerals (Rumpold and Schluter, 2013), the accessibility of certain essential minerals, such as zinc, from mealworm matrices, is rather poor (Mwangi et al., 2018). The process of drying further aggravates the issue (Kröncke et al., 2019). Zinc is an essential element, crucial for signal transduction, DNA replication, transcription, and protein synthesis, performing structural, catalytic, and regulatory functions (Hall and King, 2023; Qi et al., 2024). Additionally, zinc competes with heavy metal ions (e.g., cadmium) for transporter-mediated ion transport, lowering their cellular absorption. Zinc addition to the diet raised larval zinc levels without triggering developmental abnormalities, suggesting that the tissues, gastrointestinal tract included, were not impacted by acute toxic effects, thus shielding cells from the toxic effects of heavy metals (Keil et al., 2020; Yu et al., 2021). Beyond its availability to the insect’s gut cells, zinc might also play a key role in host–microbe communication (Hrdina and Iatsenko, 2022; Khan and Lang, 2023).

Zinc is essential for the bacterial microbiome, as zinc-binding proteins constitute 5% of the bacterial proteome (Andreini et al., 2006), with microbiota utilizing up to 20% of the dietary zinc consumed by the host (Smith et al., 1972). However, excessive amounts of zinc can lead to cytotoxic effects on bacterial cells, as zinc competes with the binding of other metal ions in active centres of enzymes, leading to the inactivation of important enzymes (Gaupp et al., 2012). The effects of dietary zinc on insect gut microbiome have been scarcely investigated, as most data on livestock are derived from studies of piglets undergoing weaning. These portray a range of variations in microbiome composition depending on the source and amount of zinc (Oh et al., 2021; Ortiz Sanjuan et al., 2022; Pieper et al., 2020; Wang et al., 2019). The application of specific diets and rearing strategies to manipulate microbial communities is showing great potential in improving TM (Guo et al., 2025; Sibinga et al., 2025). While integrating zinc in combination with probiotics in insect rearing may enhance microbiome balance and overall health, the approach is still in its early stages and demands further study to establish effective protocols.

The focus of this study was on the fermentation of agricultural side-streams, namely spinach, to naturally develop complex probiotic populations to enrich the diet of TM larvae. The idea was to formulate diets that can promote the health of insect hosts, thus resulting in a more nutritious food/feed source, and featuring a safer microbiome profile. Although probiotics are considered an alternative solution to antibiotic use, their effects on growth, development, nutritional value and reduction of the microbial load of edible insects have not been adequately studied. Moreover, the ability to include agricultural side-streams as feed (and moisture) sources can lead to more cost-effective diets, while reducing overall waste (Adamaki-Sotiraki et al., 2024; Van Peer et al., 2021). Our approach focused on evaluating microbial valorisation of agricultural side-streams through pre-fermentation and its effects on the mealworm microbiome when provided together with ZnSO_4_.

Firstly, the bacterial community composition of spinach fermented up to 96h, was assessed by 16S metabarcoding. Subsequentially, early-stage TM larvae were fed diets supplemented with fermented agricultural side-streams and ZnSO_4_ for 35 days. We investigated the effect of fermented spinach and zinc-containing diets on the survival and growth parameters of TM larvae. Additionally, accumulated zinc was quantified. Furthermore, a comprehensive nutritional analysis of the feeding materials used to compose the experimental diets was performed, featuring dry matter and protein content, presence of select organic acids, free α-amino nitrogen (FAN), free amino acids and volatile compounds. Lastly, 16S genomic sequencing of larvae fed supplemented diets was performed to access microbiota changes resulting from exposure to different treatments.

## Materials and Methods

### Spontaneous fermentation of spinach

Fresh organic spinach (REWE, Germany) was rinsed to remove surface debris and, subsequently, ground to a coarse slush using a mortar and pestle. The spinach slush was then incubated for spontaneous fermentation, without a starter culture, for 24, 48 and 96h in 50 mL falcon tubes, at 30°C and 50 rpm. After the respective fermentation periods elapsed, spinach material was frozen at -20°C until needed (including a non-fermented control; NF). Prior to freeze storing, samples from all fermentation periods were plated in LAB-selective MRS agar (Carl Roth, Karlsruhe, Germany), incubated for 24h at 37°C and formed single colonies were collected for 16S sequencing.

### Feed preparation

All the feed used was based on a standard wheat bran diet (dry feed) composed as follows: 87% wheat bran (Mehlwurmfarm GmbH, Germany), 7% yeast mix (5%, m/m, yeast extract (Sigma Aldrich, Germany) in wheat flour), 5% oat flakes (REWE, Germany) and 1% soy protein (Sigma Aldrich, Germany). Apple (REWE, Germany) was added to the standard diet as the source of moisture, as this formulation was used in regular feed for *Tenebrio molitor*, but also as the experimental control in the feeding trials. In spinach-based diets, non-fermented (NF) or 96h fermented (F96h) spinach slushes replaced the apple as the source of moisture and probiotics (10% m/m). Additionally, zinc, provided as ZnSO_4_·7H_2_O (Carl Roth, Karlsruhe, Germany), was added in varying quantities to the standard wheat bran diet and freeze-dried, prior to the addition of spinach, to formulate Zn_low_ and Zn_high_ diets. For the diet lacking zinc fortification, only double-distilled water was mixed into the wheat bran diet before freeze-drying. A summary of all diets included in the feeding trial can be found in Table S1.

### Feeding trial design

The feeding trial protocol was derived from Keil et al. (2020), with minor modifications to suit our setup. Adult *T. molitor* beetles from our lab stock, started from a population ordered from Mehlwurmfarm GmbH in 2023, were placed in standard wheat bran diet for oviposition. After 7 days, the beetles were removed from the laying substrate, leaving the eggs. Freshly hatched larvae were allowed to grow in standard wheat bran diet for 1-2 weeks. TM larvae (stage L_1–2_) were seeded into 400 ml glass beakers containing the formulated diets (Table S1). Each treatment was administered in 3 biological replicates, each comprised of 30 larvae (density of 71 larvae/dm^2^ growth surface). All beakers were incubated at 28 °C and 75% relative humidity (with no circadian rhythm for light, temperature or relative humidity) for 5 weeks. Each group was fed fresh dry feed weekly, while the wet feed (i.e., apple or spinach) was replaced twice a week. Larval weight and survival were assessed on a weekly basis until the terminus of the feeding trials (35 days), when the first pupae were found. Larvae were weighed individually, but not kept individually, therefore larval weight gain was calculated by subtracting the average initial weight to that of the subsequential weekly scaling. Moreover, survival was measured by dividing the number of live larvae at a given time point by the starting number of larvae, then multiplying the resulting value by 100, and therefore expressed as a percentage. Before further processing, TM larvae were harvested, starved for 24 h to diminish gastrointestinal feed residues prior to further analyses, and then ethically sacrificed by flash freezing at -80 °C (Delvendahl et al., 2022). All samples were later stored at -20 °C.

### DNA isolation

Genomic DNA of single colonies collected from MRS agar plates after fermentation were isolated using GeneMATRIX Gram Plus and Yeast genomic DNA purification kit (Roboklon, Germany) following manufacturer’s instructions. Protocol was adjusted by using a cell disruptor for mechanical lysis (Retsch MM 400, Thermo Scientific, Germany), instead of a vortex adapted with a tube holder, for 10 min. at 30 Hz, as suggested by manufacturer’s instructions.

Genomic DNA of feeding material and TM larvae were isolated using the DNeasy PowerSoilPro Kit (Qiagen Inc., Germany) following manufacturer’s instructions. This protocol was also adjusted by using a cell disruptor for mechanical lysis (Retsch MM 400, Thermo Scientific, Germany), instead of a vortex adapted with a tube holder, for 10 min. at 30 Hz. Additionally, upon adding solution CD1 to the samples, a 10 min. incubation at 65°C was also included to further potentiate the action of CD1.

The DNA quantity and quality were estimated by measuring the optical density at A260/280 using the Nanodrop spectrophotometer (Thermo Scientific, Germany) and the Qubit 4 Fluorometer (Thermo Scientific, Germany).

### 16S amplicon library preparation and sequencing

16S rRNA gene amplicon libraries were prepared by PCR amplification of an approximate 467 bp region within the hypervariable (V3–V4) region of the 16S rRNA gene in bacteria. The starting material consisted of 50 ng of each of the extracted and purified DNA, and nuclease-free water (non-template control), respectively, according to the Illumina 16S metagenomic sequencing library protocol, with modifications (Illumina, 2013a).

PCR was initially performed with broad spectrum 16S rRNA primers (fw: 5′-GGTGYCAGCMGCCGCGGTAA-3′, rv: 5′-CCGYCAATTYMTTTRAGTTT-3′), using Q5® Hot Start High-Fidelity 2X Master Mix (New England Biolabs, USA). Cycle conditions were 98 °C (30 s), then 30–35 cycles of 98 °C (5–10 s), 50–72 °C (10–30 s), 72 °C (30 s), and a final extension of 72 °C (2 min). Libraries were purified using NGS beads (GCbiotech, The Netherlands). Dual indices and Illumina sequencing adapters from the Illumina Nextera XT index kits v2 B and C (Illumina, San Diego, USA) were added to the target amplicons in a second PCR step using using Q5® Hot Start High-Fidelity 2X Master Mix (New England Biolabs, USA). Cycle conditions were 98 °C (30 s), then 8 cycles of 98 °C (10 s), 67 °C (30 s), 72 °C (30 s), and a final extension of 72 °C (2 min). Libraries were purified again using NGS beads.

Libraries were quantified on a Qubit Fluorometer (Thermo Scientific, Germany). The barcoded amplicon libraries were combined in equal concentrations into a single pool according to their quantification measurement. Two μl of each negative control library were added to the pool. The library pool was quantified again using the Qubit Fluorometer and the size was assessed with an Agilent DNA 1000 Kit (Agilent Technologies Ireland Ltd., Ireland) on an Agilent 2100 Bioanalyser (Agilent Technologies Ireland Ltd., Ireland).

The library pool was diluted and denatured according to the Illumina MiSeq library protocol (Illumina, 2013b). The sequencing run was conducted on the Illumina MiSeq using the 500 cycle MiSeq reagent kit to produce 250 bp paired-end reads.

### 16S bacterial population analysis

The primer sequences were removed from the raw reads using cutadapt (Martin, 2011). We checked the quality of the reads with FastQC (Andrews, 2010). 16S rRNA amplicons were then processed according to the full workflow implemented in the DADA2 R package (Callahan et al., 2016). Briefly, we trimmed the reads obtained from diet samples or larval samples according to their quality; reads obtained from the diets were trimmed at 280 bp (forward) and 200 bp (reverse), whereas reads coming from larvae were trimmed at 230 bp (forward) and 120 bp (reverse) with the ‘filterAndTrim’ function. The reads were then denoised according to their error rates with the ‘dada’ function, and merged, before a chimera removal step. The reads were then processed into Amplicon Sequence Variants (ASVs), and ASVs were assigned Operational Taxonomic Units (OTUs) with the ‘assignTaxonomy’ function and using the version 18 of the Ribosomal Database Project (RDP; (Maidak et al., 1997)). Reads identified as mitochondria or chloroplasts were excluded from further analysis.

### Zinc quantification of feeding material and mealworm larvae

We implemented the protocol outlined by Keil et al. (2020) with minimal modifications. Feed samples were ground replica-wise and portioned before heating in a laboratory microwave digester (Mars 6, CEM GmbH, Kamp-Lintfort, Germany) in a 1:1 mixture of ultrapure HNO_3_ (65%) and H_2_O_2_ (30%) containing 500 µg/L indium as an internal standard to estimate metal recovery rates. A 3:1 mixture of ultrapure HNO_3_ (65%) and H_2_O_2_ (30%) plus 500 µg/L indium was used for crashing the larvae. After digestion the samples were prediluted to 5 ml using 18.2 MΩ·cm water (Millipore Milli-Q Water Purification System). Samples were further diluted 1:100 in 0.65% HNO_3_ containing 5 µg/L rhodium and analysed on a NexION® 1000 inductively coupled plasma mass spectrometer (ICP-MS; PerkinElmer LAS GmbH, Rodgau, Germany; (Maares et al., 2025)). ICP-MS conditions are listed in Table S2. The instrument was tuned daily for maximum sensitivity with a NexION® setup solution (<0.03 oxide ratio (^140^Ce^+^ ^16^O/^140^Ce^+^), double charged ratio <0.03 (^137^Ba^++^/^137^Ba^+^) and background counts <2 cps). The reliability of our zinc analysis was verified with the reference material Fortified Breakfast Cereal (NIST-3233), which has a certified content of 628 ± 16 mg/kg, while our determined value was 677 ± 56 mg/kg.

The Bioaccumulation Factor (BAF) was calculated as previously described (Walker, 1990):

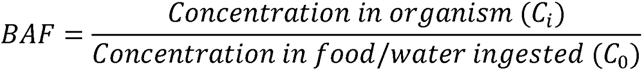

### Nutritional and anti-nutritional assessment of diet composition

The dry matter percentage of spinach (non-fermented and fermented for 24, 48 and 96h) and all formulated diets was determined using a halogen moisture analyser HR73 (Mettler Toledo, Schwerzenbach, Switzerland). The samples were heated in the weighing pan of the built-in scale by a circular halogen lamp, according to the manufacturer’s standard program.

Protein content was determined by the Kjeldahl method and calculated by multiplying the measured nitrogen content by a factor of 6.25, according to DIN EN 25663 and the Association of German Agricultural Analytic and Research Institutes (Forschungsanstalten, 2013). Results are shown as grams of protein per 100 g of dry matter (g/100 g DM).

The phytic acid content was determined using the Phytic acid/Total Phosphorus (K-PHYT) enzyme kit (MegaEnzyme, Ireland), according to manufacturer’s instructions. The kit measures phosphorus released from a ground sample after treatment with phytase and alkaline phosphatase. Samples not treated with phytase allow the quantification of monophosphates not associated with phytic acid, which is necessary to avoid an overestimation of phosphorus in phytic acid. Since phosphorus comprises 28.2% of phytic acid, multiplying phytic acid-phosphorus by a factor of 3.55 yields the amount of phytic acid. Oat samples provided with the kit were used as control. Results are shown as grams of phytic per 100 g of dry matter (g/100 g DM).

Organic acids were analysed by GC-MS after derivatisation with N-tert-butyldimethylsilyl-N-methyltrifluoroacetamide (MTBSTFA). The freeze-dried and ground samples were extracted with 6 M hydrochloric acid, in triplicates. The supernatants were combined and heated at 100 °C for 1h. The samples were then concentrated using a vacuum concentrator (SpeedVac; Jouan RC 10.22 RCT 90, Dreieich, Germany) and refrigerated (Cooling Trap; Jouan RCT90, Dreieich, Germany), until fully dry, to determine the extracted mass. The samples were redissolved in 6 M HCl, filtered and 20 µL of each sample were freeze-dried (Christ Beta 1–8, Martin Christ, Germany). To each sample, 50 µL of pyridine and 50 µL of the derivatization reagent MTBSTFA were then added. Derivatization was carried out for 4 h at 100 °C. Subsequentially, ethyl acetate was added and samples were analysed by gas chromatography-mass spectrometry (GC-MS). To identify and quantify the existing organic acids, the following standard solutions were prepared: oxalic acid, lactic acid, malic acid, citric acid and levulinic acid. The parameters used for GC-MS measurements, including the temperature program, are summarized in Table S3. A quantitative result was only possible for oxalic acid, shown as milligrams of oxalic acid per 100 g of dry matter (mg/100 g DM). Qualitative results were produced for the remaining organic acids by calculating the area of the respective peaks and shown as area per gram of dry matter (area/g DM).

Free α-amino nitrogen (FAN) was assessed according to the modified EBC-ninhydrin method (Lie, 1973). Briefly, the determination of free amino nitrogen in samples is carried out using a colour reaction of ninhydrin with amino acids. Firstly, reagent A is prepared, consisting of disodium hydrogen phosphate dihydrate (NaLHPOL·2HLO), potassium dihydrogen phosphate (KHLPOL), fructose and ninhydrin (added last and gently vortexed), dissolved in distilled water and covered with aluminium foil. Moreover, reagent B consists of potassium iodate (KIOL) dissolved in distilled water and absolute ethanol (3:2 v/v ratio). For analysis, 20 µL of each sample are pipetted into 1.5 mL Eppendorf tubes, together with 30 µL of distilled water and 50 µL of reagent A. The mixture is heated for 5 min., at 90 °C, in a thermomixer, without shaking. Afterwards, 900 µl of reagent B is added and the solution is thoroughly vortexed. Finally, 300 µL of the mixture is pipetted into a 96-well plate and the absorbance is measured at 570 nm on a plate reader (TECAN Infinite M200, Switzerland). A standard curve with glycine was used for calibration. Results are shown as grams of FAN per 100 g of dry matter (g/100 g DM).

Free amino acids were analysed by GC-MS after derivatisation with MTBSTFA. The freeze-dried and ground samples were extracted with 6 M hydrochloric acid, in triplicates. The resulting extract was evaporated using a vacuum concentrator (SpeedVac; Jouan RC 10.22 RCT 90, Germany) and refrigerated (Cooling Trap; Jouan RCT90, Germany), until fully dry, to determine the extracted mass. The evaporated extracts were then redissolved in 0.1 M hydrochloric acid. For each sample, 100 µL were removed and replaced by 100 µL of an internal standard (norleucine, 7.4 mmol/L), prior to being freeze-dried (Christ Beta 1–8, Martin Christ, Germany). The dry samples were derivatized for 4 hours at 100°C with MTBSTFA. The derivatized samples were added 500 µL of ethyl acetate and analysed by GC-MS (following the aforementioned conditions; Table S3). Standard solutions of the 20 proteinogenic amino acids were prepared and analysed in parallel. Results are shown as milligrams of amino acid per 100 g of dry matter (g/100 g DM).

Volatile compounds were analysed by GC-MS and headspace injection, which allows the detection of short chain volatile compounds in the space above a solid sample. This method is typically used for flavour compounds, as substances are broken down into short compounds during fermentation, the extent to which volatile compounds are detectable was assessed (Keil et al., 2022). For automated sample incubation and usage, the GC-MS system was equipped with a Combi-PAL-RSI autosampler (Axel Semrau GmbH & Co. KG, Germany). Samples were incubated in the agitator module incubated at 120 °C and 500 rpm for 7 min prior to GC injection. Subsequently, 1 mL of vapor space was injected into the GC-MS system. The parameters used for GC-MS measurements, including the temperature program, are summarized in Table S4. Chemical identification was confirmed by comparing retention times and mass spectra of samples with those of analytical standards, and by using the NIST (National Institute of Standards and Technology) database. The quantitation was performed in SCAN mode (mass scan 33–350 m/z), using total ion current (TIC; expressed as abundance units (AU)), and provide semiquantitative data. Data acquisition was performed using the GCMSsolution software version 4.53SP1 (Shimadzu Deutschland GmbH, Germany).

### Statistical analysis

All data was analysed using the R software (Team, 2020). Larval weight was included as a response variable in a Linear Mixed Model (LMM) according to the diet type (control or spinach, fermented or not during different amounts of time) and zinc dosage (Zn_0_, Zn_low_ or Zn_high_) as well as the interaction between them as explanatory variables, and with the experimental replicate as a random factor (‘lmer’ function of the ‘lme4’ package; (Bates et al., 2015)). Survival of the larvae was not analysed since mortality of the larvae was negligible (less than 10 % in each treatment; Fig. 2B).

The zinc content which was measured in the larvae was log-transformed and included as a response variable in a Linear Model (LM), with diet type, zinc concentration in the diets as well as the interaction between them as explanatory variables. An LM was also used to analyse dry matter content in spinach and in the diets, as well as the content of 4 organic acids, FAN, 11 free amino acids and 2 volatile compounds in spinach (‘lm’ function of the ‘lme4’ package).

The zinc and protein content in the diets and the BAF, as well as the content of malic acid, 6 free amino acids and 5 volatile compounds in spinach was assessed with a generalised linear model (GLM), fitted for gamma distribution (‘glm’ function of the ‘lme4’ package). Moreover, a GLM fitted for a negative binomial distribution (‘glm.nb’ function of the ‘MASS’ package) was used to analyse the content in levulinic acid and 2 volatile compounds in spinach.

In all the cases described above, pairwise comparisons between treatment regimens can be performed by eye, as previously reported (Cumming, 2009; Cumming and Finch, 2005). The difference between 2 selection regimes is deemed significant when the 95% confidence intervals (95CI) do not overlap more than half of their length. Effect plots for post-hoc comparison are displayed in the Supporting information (Fig. S4–S16), on effect plots (‘effects’ package (Fox, 2003; Fox and Weisberg, 2018).

The differential abundance of OTUs between samples of diets and larvae was analysed with the ‘DESeq’ function of the ‘DESeq2’ package (Love et al., 2014). The counts of member of the genus Staphylococcus and Lactococcus, which were significantly differentially abundant between larval treatments were further analysed with a negative binomial Generalised Linear Model (GLM, ‘MASS’ package) with OTU counts as a response variable, according to the diet fermentation status and the zinc concentration as explanatory variables.

A Bray-Curtis dissimilarity matrix was used to assess the Beta-diversity among the samples obtained from larvae, and whether the distance between samples of the various treatments was significant (‘adonis2’ function of the ‘vegan’ package; (Oksanen et al., 2025)). Contrasts were performed with the ‘pairwise.adonis2’ function of the ‘pairwiseAdonis’ package (Martinez Arbizu, 2020). In the case of the samples obtained from the diets, the sample size was too low to perform an analysis, but showed clear separation on the Bray-Curtis PcoA.

The Shannon index of each sample was calculated as a measure of Alpha diversity (‘estimate_richness’ function of the ‘phyloseq’ package (McMurdie and Holmes, 2013)). It was then included as an explanatory variable in a GLM with a Gamma family (log link), according to the fermentation status and zinc concentration of the diet.

The figures of the main text and supplementary material were produced using GraphPad Prism (version 10 for Windows, GraphPad Software, Boston, Massachusetts, USA) and the ‘ggplot2’ package (Wickham, 2016).

## Results

### Spinach fermentation potentiates probiotic populations

To evaluate which length of fermentation favoured the growth of probiotic populations in spinach slushes, these were incubated for three time periods, 24h, 48h and 96h. A 16S metabarcoding analysis detected 69 genera in all the spinach/substrate samples, of which ten bacterial genera were significantly differentially abundant, at a threshold of alpha < 0.05. No further analyses were prepared due to low sample size, referring instead to Fig. 1 and Fig. S1 (corresponding to the stacked barplot and differential abundance counts, respectively).

**Figure 1.**
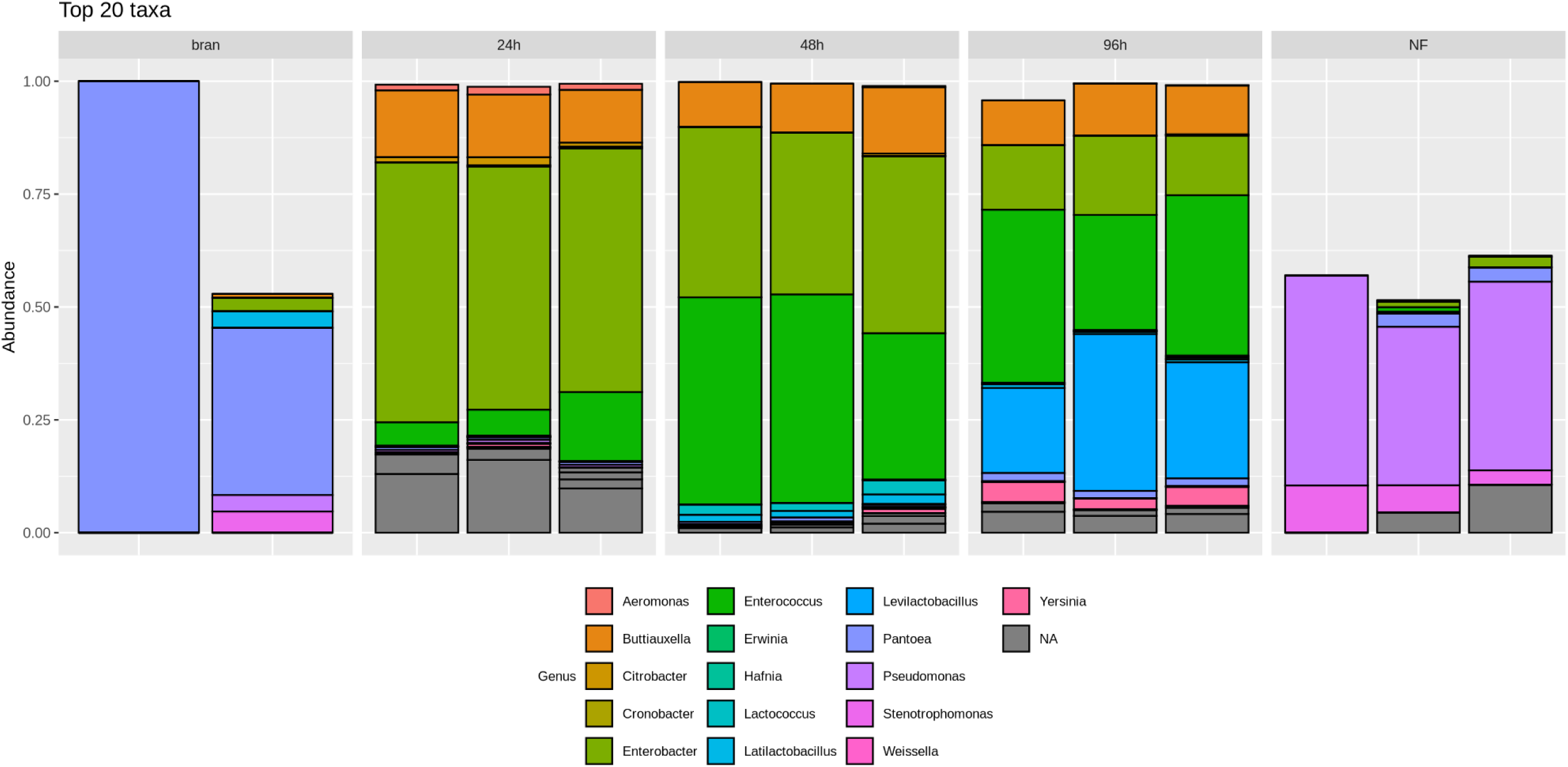
– Stacked barplot representing the relative abundance of the 20 most abundant bacterial genera detected in the different substrate and spinach samples. Each panel contains the bars corresponding to a treatment group: wheat bran (substrate), and spinach either non-fermented (NF) or fermented for 24, 48 or 96 hours. Each bar represents an individual sample, with each different coloured box representing one bacterial genus. Grey boxes indicate OTUs for which no genus could be assigned.

Fermentation at these 3 time points resulted in proliferation of bacteria from the *Enterobacteriacae* family (*Enterobacter*, *Buttiauxella*, and a taxon of unassigned genus) compared to control and non-fermented spinach. A member of the genus *Enterococcus* (family *Enterococcaceae*) followed the same pattern. This result was confirmed by plating spinach samples on MRS medium, from which small colonies were recovered, for metabarcoding. They were identified as members of the *Enterococcus* genus (Fig. S2). Fermentation also resulted in the proliferation of two members of the *Lactobacillaceae* family. Members of the genus *Latilactobacillus* were absent in non-fermented spinach but present after 24 hours of fermentation, increasing in abundance over 48 and 96 hours. *Levilactobacillus* was more abundant in spinach samples having fermented for 96 hours compared to 24 h, 48 h, non-fermented and control. Additionally, members of the *Lactococcus* genus (family *Streptococcaceae*) had higher counts at 24 and 48 hours of fermentation, but declined at 96 hours.

One member each of the *Erwiniaceae* and *Yersiniaceae* families, with unassigned genera, also proliferated from 24 hours of fermentation compared to non-fermented and control. Conversely, the prevalence of a member of the *Pseudomonas* genus (family *Pseudomonadacae*), which was present in non-fermented spinach, was significantly decreased from 48 hours of fermentation compared to non-fermented spinach, and absent after 96 hours of fermentation (Fig. 1).

A PcoA plot of the Bray-Curtis dissimilarity between samples (beta-diversity) showed a clear separation between substrate (wheat bran) samples, non-fermented and fermented spinach samples (Fig. S3). Fermented spinach samples were distinct from non-fermented samples, but the fermentation time did not separate the samples.

Taking these findings into account, 96h-fermented spinach was chosen to be used in diet formulation for feeding trials.

### Nutritional profile of fermented spinach and formulated diets

Dry matter content was found to be generally consistent among spinach samples, except for 96h-fermented spinach which showed a slight, but significant, higher percentage (Table 1 and Fig. S4; F_3,8_ = 7.964, p = 0.0087), when compared to the remaining spinach groups. Moreover, diets formulated with zinc also led to significantly higher dry matter content (except for NF + Zn_high_, which increase was not statistically significant; Table 1 and Fig. S4; F_3,8_ = 12.45, p = 0.0002), when compared to control and non-zinc supplemented diets.

**Table 1.** – Dry matter (%; DM) assessment of spinach samples and all formulated diets (as summarised in Table S1), as well as protein content of all formulated diets (as summarised in Table S1; g/100g DM). Different letters (i.e., ^a^, ^b^ and ^c^) represent statistically significant differences; * Represented by only 1 replicate.

| **Dry matter - Spinach** | **NF** | **F 24h** | **F 48h** | **F 96h** |  |  |  |
| --- | --- | --- | --- | --- | --- | --- | --- |
| Mean | 7.867<sup>a</sup> | 7.633<sup>a</sup> | 6.967<sup>a</sup> | 9.433<sup>b</sup> |  |  |  |
| Lower 95% CI | 5.009 | 6.916 | 5.793 | 9.146 |  |  |  |
| Upper 95% CI | 10.72 | 8.35 | 8.141 | 9.72 |  |  |  |
| **Dry matter - Diets** | **Control** | **NF** | **NF + Zn<sub>low</sub>** | **NF + Zn<sub>high</sub>** | **F** | **F + Zn<sub>low</sub>** | **F + Zn<sub>high</sub>** |
| Mean | 77.97<sup>a</sup> | 80.67<sup>a</sup> | 95.63<sup>b</sup> | 82<sup>a</sup> | 79.9<sup>a,*</sup> | 88.7<sup>c</sup> | 84.13<sup>c</sup> |
| Lower 95% CI | 67.76 | 69.83 | 94.76 | 74.44 | — | 82.35 | 83.62 |
| Upper 95% CI | 88.18 | 91.5 | 96.51 | 89.56 | — | 95.05 | 84.65 |
| **Protein** |  |  |  |  |  |  |  |
| Mean | 20.1<sup>a</sup> | 22.47<sup>b</sup> | 21.13<sup>a,b</sup> | 19.27<sup>a</sup> | 20.33<sup>a</sup> | 18.43<sup>a</sup> | 19.13<sup>a</sup> |
| Lower 95% CI | 17.33 | 20.35 | 20.75 | 17.99 | 20.19 | 13.39 | 18.51 |
| Upper 95% CI | 22.87 | 24.58 | 21.51 | 20.54 | 20.48 | 23.48 | 19.76 |

The level of six organic acids in spinach was assessed throughout the length of fermentation, namely citric, malic, lactic, levulinic, oxalic and phytic acid. Citric and levulinic acid showed a similar response to fermentation, being steadily degraded by the present bacterial populations after 24 and 48h, respectively (Table 2 and Fig. S6; citric acid: F_3,7_ = 24.7, p = 0.0002; levulinic acid: X^2^_3,7_ = 1890.5, p < 0.0001). Despite displaying a similar trend, after being significantly degraded at the early stage of fermentation, the malic acid content rebounds in later stages, as it’s level after 96h was not significantly different from that of non-fermented spinach (X^2^_3,7_ = 25.765, p < 0.0001). Lactic acid content of all fermented spinach was significantly higher than that of its non-fermented counterpart, peaking at 48h, then experiencing some degradation at 96h (F_3,7_ = 50.9, p < 0.0001). Similarly, phytic acid assessment revealed a progressive increase in concentration over fermentation time, herein significantly peaking at 96h (F_3,7_ = 5.935, p = 0.0245). Despite, no phytic acid content was detected in any of the formulated diets (wheat bran base plus source of moisture). Finally, we observed an increase in oxalic acid as a result of fermentation as well, however no statistically significant differences were found between any of the tested groups (F_3,8_ = 1.173, p = 0.386).

**Table 2.**
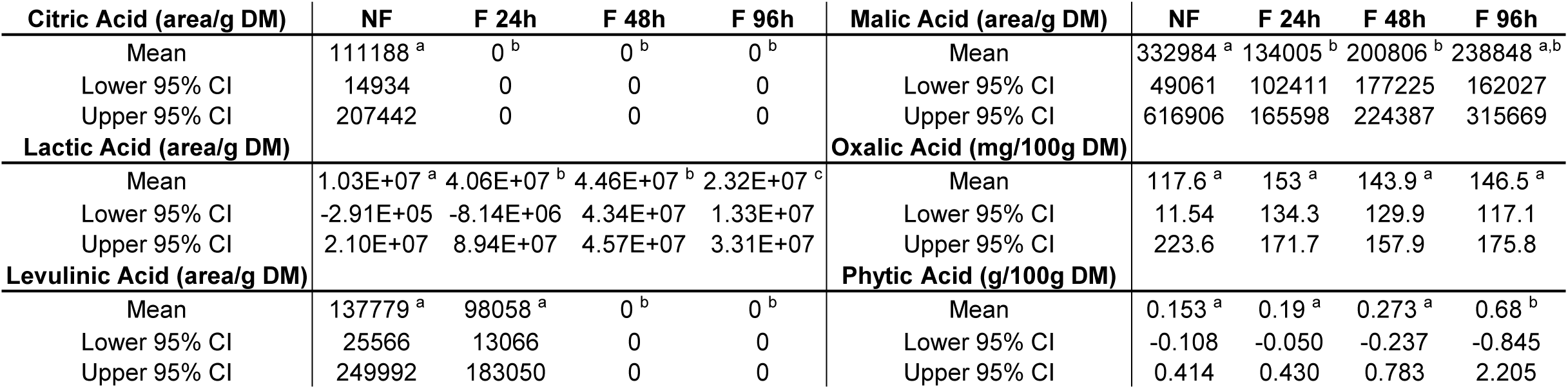
– Organic acids assessment of spinach samples. Different letters (i.e., ^a^, ^b^ and ^c^) represent statistically significant differences.

| **Citric Acid (area/g DM)** | **NF** | **F 24h** | **F 48h** | **F 96h** | **Malic Acid (area/g DM)** | **NF** | **F 24h** | **F 48h** | **F 96h** |
| --- | --- | --- | --- | --- | --- | --- | --- | --- | --- |
| Mean | 111188 <sup>a</sup> | 0 <sup>b</sup> | 0 <sup>b</sup> | 0 <sup>b</sup> | Mean | 332984 <sup>a</sup> | 134005 <sup>b</sup> | 200806 <sup>b</sup> | 238848 <sup>a,b</sup> |
| Lower 95% CI | 14934 | 0 | 0 | 0 | Lower 95% CI | 49061 | 102411 | 177225 | 162027 |
| Upper 95% CI | 207442 | 0 | 0 | 0 | Upper 95% CI | 616906 | 165598 | 224387 | 315669 |
| **Lactic Acid (area/g DM)** |  |  |  |  | **Oxalic Acid (mg/100g DM)** |  |  |  |  |
| Mean | 1.03E+07 <sup>a</sup> | 4.06E+07 <sup>b</sup> | 4.46E+07 <sup>b</sup> | 2.32E+07 <sup>c</sup> | Mean | 117.6 <sup>a</sup> | 153 <sup>a</sup> | 143.9 <sup>a</sup> | 146.5 <sup>a</sup> |
| Lower 95% CI | -2.91E+05 | -8.14E+06 | 4.34E+07 | 1.33E+07 | Lower 95% CI | 11.54 | 134.3 | 129.9 | 117.1 |
| Upper 95% CI | 2.10E+07 | 8.94E+07 | 4.57E+07 | 3.31E+07 | Upper 95% CI | 223.6 | 171.7 | 157.9 | 175.8 |
| **Levulinic Acid (area/g DM)** |  |  |  |  | **Phytic Acid (g/100g DM)** |  |  |  |  |
| Mean | 137779 <sup>a</sup> | 98058 <sup>a</sup> | 0 <sup>b</sup> | 0 <sup>b</sup> | Mean | 0.153 <sup>a</sup> | 0.19 <sup>a</sup> | 0.273 <sup>a</sup> | 0.68 <sup>b</sup> |
| Lower 95% CI | 25566 | 13066 | 0 | 0 | Lower 95% CI | -0.108 | -0.050 | -0.237 | -0.845 |
| Upper 95% CI | 249992 | 183050 | 0 | 0 | Upper 95% CI | 0.414 | 0.430 | 0.783 | 2.205 |

As expected, the protein content of the formulated diets was the highest in the diet containing non-fermented spinach and in the absence of zinc and a low zinc concentration. It was similar throughout the remaining diets (Table 1 and Fig. S5; X^2^ = 32.231, p < 0.0001).

Concomitantly, the levels of free α-amino nitrogen (FAN) significantly increased during the fermentation, doubling the level at 48h in comparison to non-fermented spinach. Nonetheless, at 96h FAN content fell back to the initial level (Table 3 and Fig. S7; F_3,4_ = 186.3, p < 0.0001).

**Table 3.** – Free amino nitrogen (FAN; g/100 g DM) assessment of spinach samples. Different letters (i.e., ^a^, ^b^ and ^c^) represent statistically significant differences.

| **FAN** | **NF** | **F 24h** | **F 48h** | **F 96h** |
| --- | --- | --- | --- | --- |
| Mean | 0.441 <sup>a</sup> | 0.722 <sup>b</sup> | 0.888 <sup>c</sup> | 0.460 <sup>a</sup> |
| Lower 95% CI | 0.258 | 0.52 | 0.84 | 0.169 |
| Upper 95% CI | 0.623 | 0.923 | 0.936 | 0.752 |

GC-MS analysis revealed that all proteinogenic amino acids, except for cysteine, asparagine and glutamine, were detected in spinach samples. Results show a clear tendency regarding abundancy, as the great majority of present amino acids peaked at 24h fermentations (Table 4, Fig. S8, S9 and S10; alanine: F_3,4_ = 9.393, p = 0.0277; arginine: X^2^_3,4_ = 266.72, p < 0.0001; aspartic acid: X^2^_3,4_ = 219.78, p < 0.0001; glutamic acid: F_3,4_ = 9.26, p = 0.0284; histidine: X^2^_3,4_ = 369.43, p < 0.0001; isoleucine: F_3,4_ = 13.42, p = 0.0148; leucine: F_3,4_ = 11.27, p = 0.0202; lysine: F_3,4_ = 26.09, p = 0.0044; methionine: X^2^ = 220.06, p < 0.0001; phenylalanine: F_3,4_ = 11.12, p = 0.0207; proline: X^2^_3,4_ = 105.26, p < 0.0001; serine: X^2^_3,4_ = 98.003, p < 0.0001; threonine: F_3,4_ = 10.39, p = 0.0233; valine: F_3,4_ = 11.59, p = 0.0193). Noteworthy, the abundance of aspartic acid, histidine, lysine and serine in 24h-fermented spinach, although higher, was not significantly different of that of non-fermented samples. Similarly, the abundance of alanine, glutamic acid, isoleucine, leucine, methionine, phenylalanine, threonine and valine in 24h-fermented spinach was not significantly different of that of 48h-fermented samples. Some exceptions to the above-mentioned trend could be found, in glycine, which peaked at a 48h fermentation (although not significantly different to that of 24h-fermented samples; Table 4 and Fig. S8; F_3,4_ = 14.9, p = 0.0123), tryptophan and tyrosine, which were only detected in non-fermented spinach (Table 4 and Fig. S10; tryptophan: F_3,4_ = 1418, p < 0.0001; tyrosine: F_3,4_ = 88.18, p = 0.0004). Interestingly, in 96h-fermented samples, the level of all detected amino acids significantly regressed to that of non-fermented spinach, or even below that.

**Table 4.** – Free amino acids (mg/100 g DM) assessment of spinach samples. Different letters (i.e., ^a^, ^b^ and ^c^) represent statistically significant differences.

| **Alanine** | **NF** | **F 24h** | **F 48h** | **F 96h** | **Methionine** | **NF** | **F 24h** | **F 48h** | **F 96h** |
| --- | --- | --- | --- | --- | --- | --- | --- | --- | --- |
| Mean | 2.856<sup>a</sup> | 130.459<sup>b</sup> | 113.024<sup>b</sup> | 68.805<sup>a,b</sup> | Mean | 2.009<sup>a</sup> | 48.038<sup>b</sup> | 20.924<sup>b</sup> | 0<sup>c</sup> |
| Lower 95% CI | -3.395 | -323.655 | -1.567 | 13.330 | Lower 95% CI | -0.692 | -120.974 | -43.700 | 0.000 |
| Upper 95% CI | 9.107 | 584.572 | 227.614 | 124.280 | Upper 95% CI | 4.709 | 217.049 | 85.548 | 0.000 |
| **Arginine** |  |  |  |  | **Phenylalanine** |  |  |  |  |
| Mean | 27.107<sup>a</sup> | 88.108<sup>b</sup> | 0<sup>c</sup> | 2.186<sup>d</sup> | Mean | 14.134<sup>a</sup> | 361.434<sup>b</sup> | 245.986<sup>a,b</sup> | 2.522<sup>a</sup> |
| Lower 95% CI | -28.838 | -294.031 | 0.000 | 0.610 | Lower 95% CI | -21.133 | -775.022 | -479.615 | -26.442 |
| Upper 95% CI | 83.052 | 470.247 | 0.000 | 3.762 | Upper 95% CI | 49.400 | 1497.890 | 971.587 | 31.485 |
| **Aspartic Acid** |  |  |  |  | **Proline** |  |  |  |  |
| Mean | 137.188<sup>a</sup> | 188.211<sup>a</sup> | 21.423<sup>b</sup> | 1.63<sup>c</sup> | Mean | 110.857<sup>a</sup> | 1787.928<sup>b</sup> | 515.41<sup>c</sup> | 40.527<sup>a</sup> |
| Lower 95% CI | -109.185 | -492.105 | -35.831 | 0.004 | Lower 95% CI | -160.344 | -3943.797 | -2043.683 | -22.858 |
| Upper 95% CI | 383.561 | 868.527 | 78.677 | 3.256 | Upper 95% CI | 382.058 | 7519.652 | 3074.503 | 103.911 |
| **Glutamic Acid** |  |  |  |  | **Serine** |  |  |  |  |
| Mean | 151.149<sup>a</sup> | 885.83<sup>b</sup> | 777.596<sup>b</sup> | 12.389<sup>a</sup> | Mean | 8.792<sup>a</sup> | 30.445<sup>a</sup> | 2.145<sup>b</sup> | 0.359<sup>c</sup> |
| Lower 95% CI | -169.009 | -2726.925 | 248.611 | 9.231 | Lower 95% CI | -10.363 | -54.973 | -10.187 | -0.639 |
| Upper 95% CI | 471.307 | 4498.585 | 1306.581 | 15.546 | Upper 95% CI | 27.946 | 115.862 | 14.476 | 1.356 |
| **Glycine** |  |  |  |  | **Threonine** |  |  |  |  |
| Mean | 6.457<sup>a</sup> | 384.526<sup>b</sup> | 460.101<sup>b</sup> | 10.909<sup>a</sup> | Mean | 11.374<sup>a</sup> | 46.514<sup>b</sup> | 9.591<sup>a</sup> | 2.765<sup>a</sup> |
| Lower 95% CI | -12.012 | -806.840 | -585.416 | -0.025 | Lower 95% CI | -20.017 | -98.883 | -25.110 | -23.442 |
| Upper 95% CI | 24.925 | 1575.891 | 1505.618 | 21.842 | Upper 95% CI | 42.764 | 191.911 | 44.292 | 28.971 |
| **Histidine** |  |  |  |  | **Tryptophan** |  |  |  |  |
| Mean | 0.494<sup>a</sup> | 0.648<sup>a</sup> | 0.223<sup>b</sup> | 0.014<sup>c</sup> | Mean | 241.339<sup>a</sup> | 0<sup>b</sup> | 0<sup>b</sup> | 0<sup>b</sup> |
| Lower 95% CI | 0.418 | -0.706 | -0.057 | -0.011 | Lower 95% CI | 159.898 | 0 | 0 | 0 |
| Upper 95% CI | 0.570 | 2.001 | 0.503 | 0.039 | Upper 95% CI | 322.779 | 0 | 0 | 0 |
| **Isoleucine** |  |  |  |  | **Tyrosine** |  |  |  |  |
| Mean | 9.272<sup>a</sup> | 232.499<sup>b</sup> | 196.286<sup>b</sup> | 4.063<sup>a</sup> | Mean | 15.954<sup>a</sup> | 0<sup>b</sup> | 0<sup>b</sup> | 0<sup>b</sup> |
| Lower 95% CI | -16.750 | -514.074 | -184.189 | 2.195 | Lower 95% CI | -5.634 | 0 | 0 | 0 |
| Upper 95% CI | 35.294 | 979.071 | 576.761 | 5.931 | Upper 95% CI | 37.542 | 0 | 0 | 0 |
| **Leucine** |  |  |  |  | **Valine** |  |  |  |  |
| Mean | 0.653<sup>a</sup> | 31.803<sup>b</sup> | 28.569<sup>b</sup> | 0.338<sup>a</sup> | Mean | 12.392<sup>a</sup> | 377.124<sup>b</sup> | 343.123<sup>b</sup> | 6.777<sup>a</sup> |
| Lower 95% CI | -1.107 | -92.578 | -9.696 | 0.160 | Lower 95% CI | -22.709 | -945.211 | -393.595 | 4.007 |
| Upper 95% CI | 2.412 | 156.184 | 66.833 | 0.516 | Upper 95% CI | 47.492 | 1699.459 | 1079.841 | 9.547 |
| **Lysine** |  |  |  |  |  |  |  |  |  |
| Mean | 95.313<sup>a</sup> | 96.427<sup>a</sup> | 28.29<sup>b</sup> | 6.487<sup>b</sup> |  |  |  |  |  |
| Lower 95% CI | -98.990 | 88.707 | -93.486 | -5.584 |  |  |  |  |  |
| Upper 95% CI | 289.616 | 104.146 | 150.066 | 18.558 |  |  |  |  |  |

Conversely to free amino acid abundancy, 96h-fermented spinach registered the highest levels of most of the detected volatile compounds, analysed by GC-MS (Table 5, Fig. S11 and S12; 2-Ethylfuran: X^2^_3,8_ = 372.63, p < 0.0001; 2-Methylbutanal: X^2^_3,8_ = 478.7, p < 0.0001; 2-Methyl-1-butanol: X^2^_3,8_ = 3066.6, p < 0.0001; 3-Methylbutanal: F_3,8_ = 6.605, p = 0.0148; Benzaldehyde: F_3,8_ = 5.679, p = 0.0221; Dimethyl disulfide: X^2^_3,8_ = 508.87, p < 0.0001; Hexanal: X^2^_3,8_ = 13236, p < 0.0001). One should point out that, 2-Ethylfuran, 2-Methylbutanal and 3-Methylbutanal abundance in 96h-fermented spinach, despite higher, was not significantly different of 24h- and/or 48h-fermented samples. Additionally, no volatile compounds were detected in non-fermented spinach. Some exceptions to the most common trend were also found, where the highest abundance for Ethyl acetate was registered in 24h-fermented spinach (Table 5 and Fig. S12; Ethyl acetate: X^2^_3,8_ = 1545.3, p < 0.0001), whereas 2-Butanone peaked in 48h-fermented samples (Table 5 and Fig. S11; X^2^_3,8_ = 631.13, p < 0.0001). However, only in Ethyl acetate abundancy in 24h-fermented spinach was found to be statistically different from that of 96h-fermented samples.

**Table 5.** – Volatile compounds (AU) assessment of spinach samples. Different letters (i.e., ^a^, ^b^ and ^c^) represent statistically significant differences.

| **2-Butanone** | **NF** | **F 24h** | **F 48h** | **F 96h** | **Benzaldehyde** | **NF** | **F 24h** | **F 48h** | **F 96h** |
| --- | --- | --- | --- | --- | --- | --- | --- | --- | --- |
| Mean | 0<sup>a</sup> | 26723<sup>b</sup> | 49838<sup>c</sup> | 33107<sup>b,c</sup> | Mean | 0<sup>a</sup> | 45595<sup>a</sup> | 40169<sup>a</sup> | 118823<sup>b</sup> |
| Lower 95% CI | 0 | 12357 | -14784 | 16575 | Lower 95% CI | 0 | 12561 | -15680 | -47689 |
| Upper 95% CI | 0 | 41090 | 114460 | 49639 | Upper 95% CI | 0 | 78630 | 96018 | 285335 |
| **2-Ethylfuran** |  |  |  |  | **Ethyl acetate** |  |  |  |  |
| Mean | 0<sup>a</sup> | 119600<sup>b</sup> | 108123<sup>b</sup> | 115224<sup>b</sup> | Mean | 0<sup>a</sup> | 87000<sup>b</sup> | 94978<sup>b</sup> | 0<sup>a</sup> |
| Lower 95% CI | 0 | 51158 | -101183 | 151278 | Lower 95% CI | 0 | 80598 | -14389 | 0 |
| Upper 95% CI | 0 | 188042 | 317430 | 79171 | Upper 95% CI | 0 | 93402 | 204344 | 0 |
| **2-Methylbutanal** |  |  |  |  | **Dimethyl disulfide** |  |  |  |  |
| Mean | 0<sup>a</sup> | 596539<sup>b</sup> | 724913<sup>b</sup> | 1201309<sup>b</sup> | Mean | 0<sup>a</sup> | 0<sup>a</sup> | 0<sup>a</sup> | 73721<sup>b</sup> |
| Lower 95% CI | 0 | -23965 | -325929 | 179712 | Lower 95% CI | 0 | 0 | 0 | -63730 |
| Upper 95% CI | 0 | 1217042 | 1775753 | 2222905 | Upper 95% CI | 0 | 0 | 0 | 211173 |
| **3-Methylbutanal** |  |  |  |  | **Hexanal** |  |  |  |  |
| Mean | 0<sup>a</sup> | 1848824<sup>a</sup> | 2184010<sup>b</sup> | 3268963<sup>b</sup> | Mean | 0<sup>a</sup> | 45967<sup>b</sup> | 40270<sup>b</sup> | 0<sup>a</sup> |
| Lower 95% CI | 0 | 9077 | -962049 | 541051 | Lower 95% CI | 0 | 11927 | -31895 | 0 |
| Upper 95% CI | 0 | 3688571 | 5330070 | 5996875 | Upper 95% CI | 0 | 80006 | 112435 | 0 |
| **3-Methyl-1-butanol** |  |  |  |  |  |  |  |  |  |
| Mean | 0<sup>a</sup> | 30435<sup>b</sup> | 53772<sup>c</sup> | 555493<sup>d</sup> |  |  |  |  |  |
| Lower 95% CI | 0 | 14221 | 15681 | 368130 |  |  |  |  |  |
| Upper 95% CI | 0 | 46648 | 91862 | 742557 |  |  |  |  |  |

### Formulated diets did not affect mealworm survival, but affected mealworm weight

The different diets fed to TM larvae did not affect the percentage of survival, as there were almost no dead larvae recorded over the course of the feeding trial. We recorded only 2 deaths in all treatment combinations, except in the treatments non-fermented spinach without zinc fortification, where there were 3, and in fermented spinach with high-dose Zn-fortification, where there were none (Fig. 2B). Thus, these deaths were considered negligible, concluding that zinc and spinach had no lethal effect on TM larvae.

While the fermentation status of the spinach in the diet did not affect the effect of zinc on larvae weight of larvae before pupation (diet composition * zinc concentration: X² = 0.58, p = 0.7483), both diet composition and zinc concentration, as simple effects, affected the final larval weight. Larvae fed on diets containing both non-fermented and fermented spinach reached a lower final weight than control larvae (Fig. 2A and Fig. S13; diet composition: X² = 14.518, p = 0.0007039). While a Zn_low_ dose fortification did not affect larval weight gain compared to control, Zn_high_ supplementation significantly decreased it (Fig. 2A and Fig. S13; X² = 21.699, p < 0.0001).

**Figure 2.**
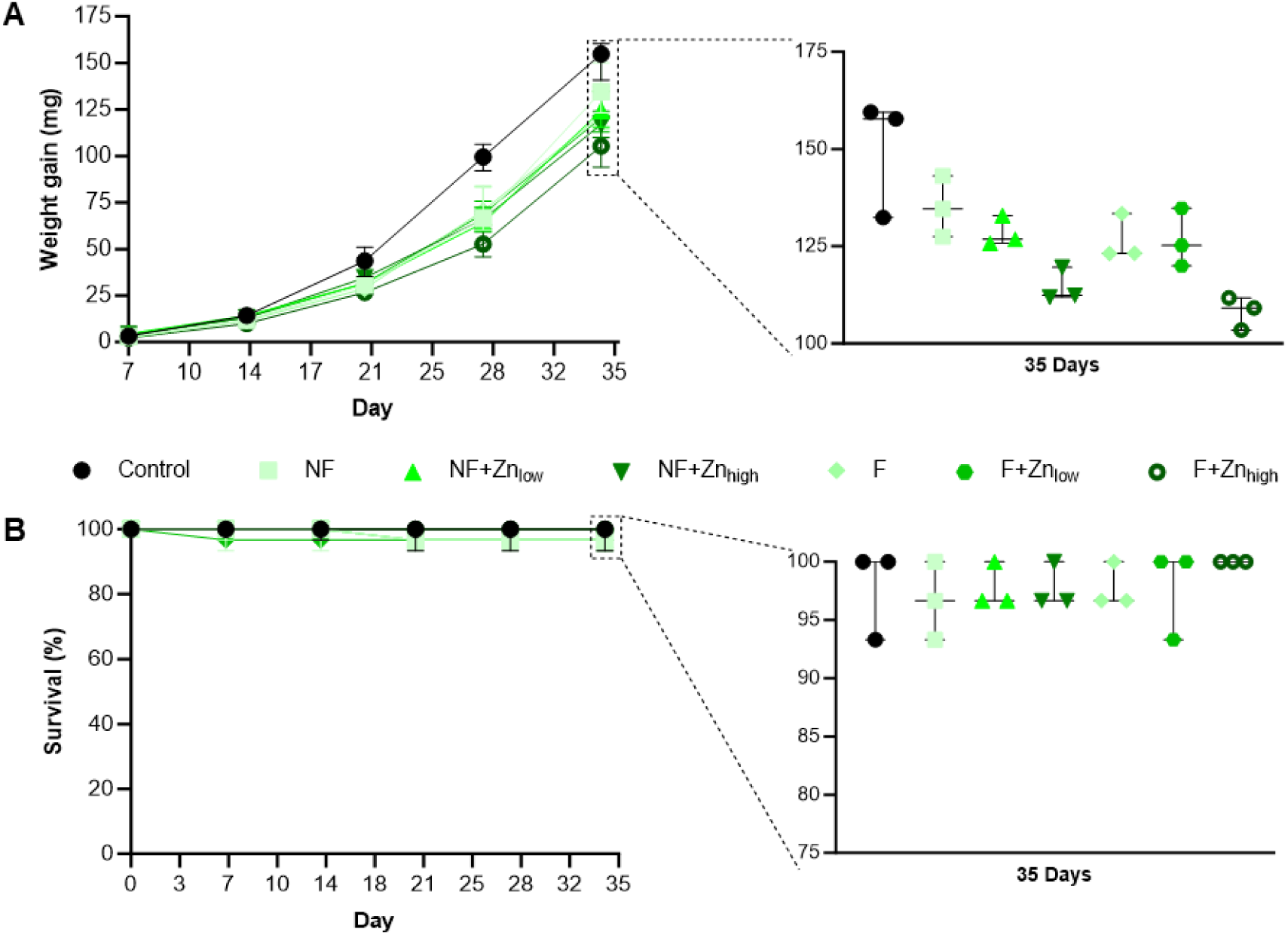
– Represents (A) Larval weight gain (mg) and (B) survival (%) throughout a 35-day feeding trial, highlighting the differences between experimental groups at the end point.

### Higher zinc supplementation led to successful accumulation in mealworm

Analysis of zinc content revealed that spinach diets without fortification were comparable to the control, while fortification resulted in significant and effective zinc incorporation (Fig. 3A and Fig. S14; X^2^_6,14_ = 221.23, p < 0.0001), peaking in the diet fortified with the highest dose (i.e., Zn_high_).

**Figure 3.**
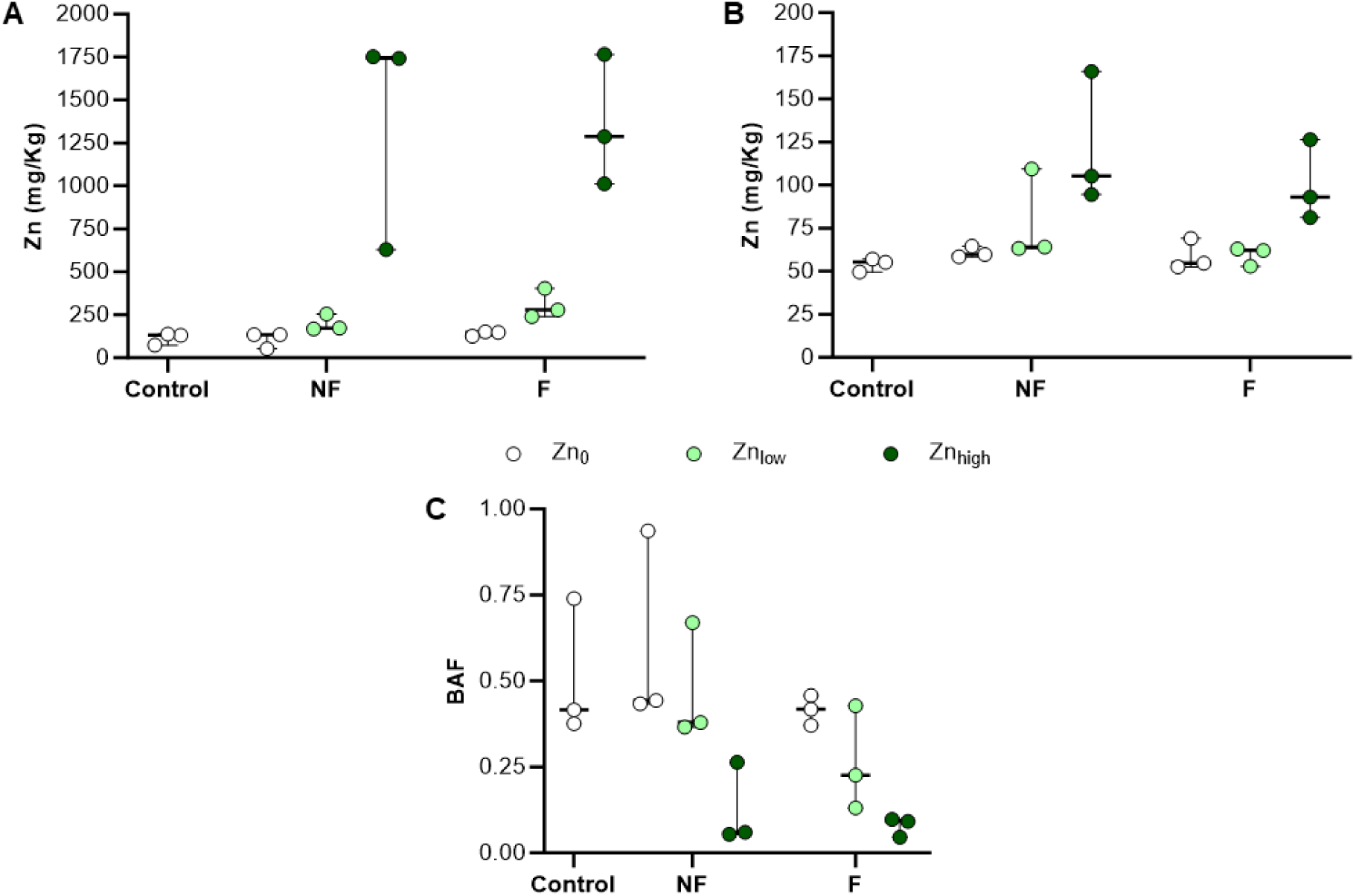
– Represents (A) zinc accumulation in the formulated diets, (B) xinc accumulation in larvae and (C) bioaccumulation factor (BAF) after 35-day feeding trial. The diets feature non-fermented (NF) and fermented (F) spinach, without zinc supplementation (Zn_0_), or fortified with low (Zn_low_) and high dosages of zinc (Zn_high_).

The fermentation status of the spinach did not significantly interfere with zinc accumulation in the larvae, according to zinc concentration in the diet (diet composition * zinc concentration: F_6,14_= 0.4398, p = 0.6527887). Similarly, the fermentation status of the diet alone did not affect the concentration of zinc found in the larvae (F_6,16_= 2.7589, p = 0.1189355). As expected, higher dietary zinc concentrations corresponded to increased zinc concentrations in the larvae (Fig. 3B and S15; F_6,18_ = 12.773, p = 0.000577). While the concentration of zinc found in larvae fed with Zn_low_ diets did not differ from that of control and larvae fed with spinach without any further zinc fortification, larvae fed Zn_high_ diets contained a significantly higher zinc concentration than that of both the other treatments.

The calculated BAF was shown to be significantly lower for diets featuring the highest zinc supplementation when compared to the remaining treatments (Fig. 3C and Fig. S16; X^2^_6,14_ = 32.66, p < 0.0001). No significant differences in BAF were found between treatments featuring low zinc supplementation, or non-fortified samples, and the control group.

### Spinach supplementation led to a mealworm microbiome overhaul

Twenty-three genera were detected in TM larvae under our experimental conditions. The consumption by the larvae of the diets containing fermented or non-fermented spinach, and zinc at different concentrations, did significantly affect the abundance of 5 bacterial taxa (Fig. 4 and Fig. S17). *Staphylococcus* was present in larvae reared on all diets, but more abundant in larvae fed spinach regardless of fermentation status or zinc concentration (diet status * zinc concentration: deviance = 2.6542, p = 0.265252, df = 6,56; zinc: deviance = 0.1266; p = 0.938676, df = 3,58; diet: deviance = 15.8864, p = 0.0003551, df = 3,60). Unlike members of the genus *Staphylococcus* which were detected in all larvae, members of the 4 remaining genera showed no counts in some of the samples.

**Figure 4.**
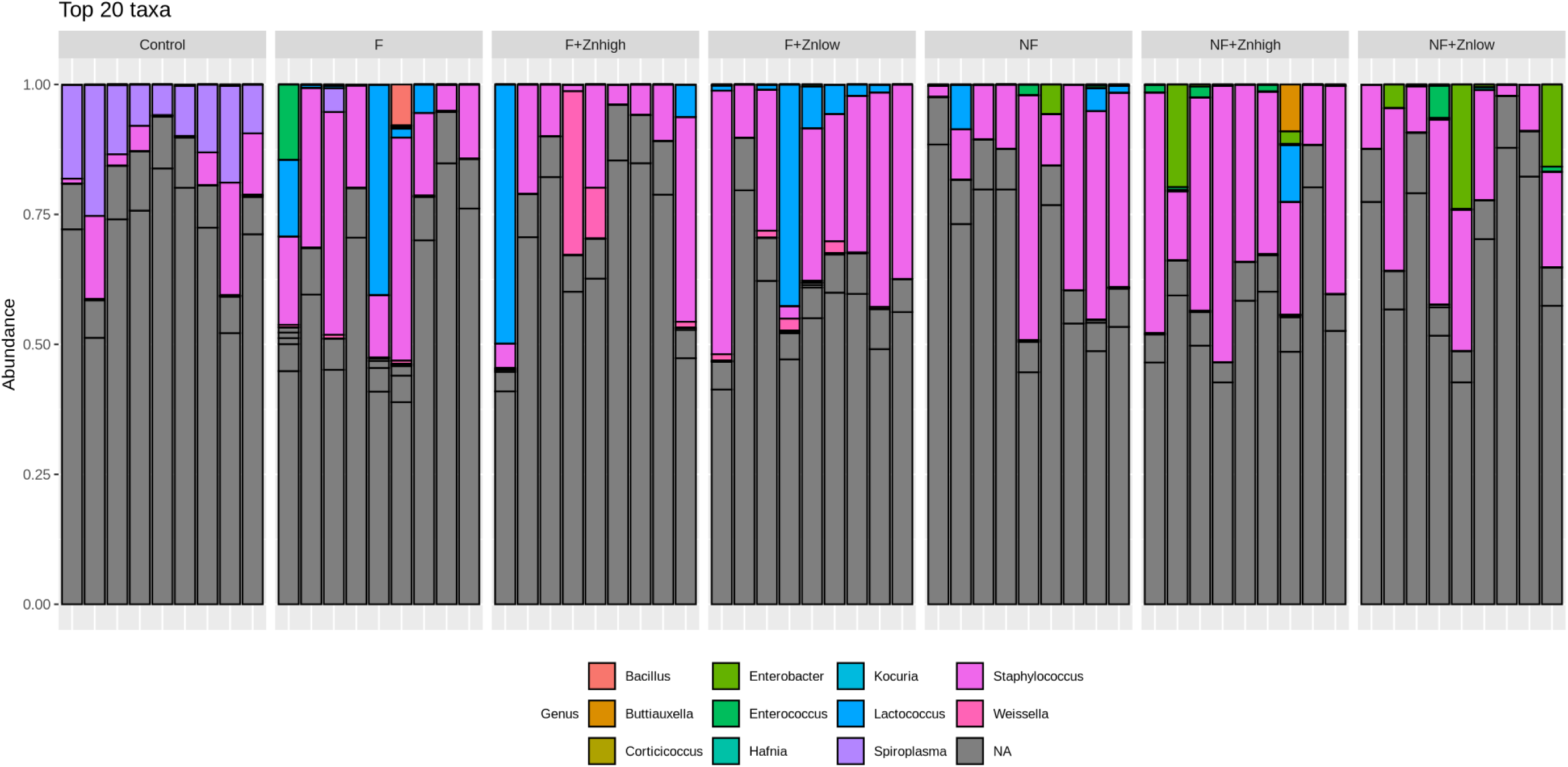
– Stacked barplot representing the relative abundance of the 20 most abundant bacterial genera detected in *T. molitor* larvae having been fed a control diet or fed diets containing spinach, either fresh or fermented for 96 hours, as well as 3 different concentrations of zinc (Zn_0_, Zn_low_ or Zn_high_). Each bar represents an individual sample, with each different coloured box representing one bacterial genus. Grey boxes indicate OTUs for which no genus could be assigned.

The counts of the genus *Lactococcus* were not explained by the concentration of zinc in the diets, neither by interaction with the diet (diet * zinc concentration: deviance = 53.323, p < 0.0001, df = 6,56), nor as a simple effect (zinc concentration: deviance = 0.57529, p = 0.75, df = 3,58). Counts of *Lactococcus* were however explained by the diet (deviance = 10.765, p = 0.004597, df = 3,60), as more counts were found in larvae reared on fermented than that of non-fermented spinach. Moreover, both these treatments had higher counts than the control (diet: deviance = 10.765, p = 0.004597, df = 3,60), since most control larvae did not have any counts of *Lactococcus*.

The counts of the genus *Lactococcus* were explained by the interaction between zinc and fermentation status (diet * zinc concentration: deviance = 53.323, p < 0.0001, df = 6,56). Most control larvae did not have any counts of *Lactococcus*. The same pattern was observable in F+Zn_high_ and NF+Zn_low_ larvae. In fermented spinach, counts of Lactococcus increased at Zn_0_ and Zn_low_ compared to control, but collapse at Zn_high_. In non-fermented spinach however, while Zn_0_ has similar levels to that of F+Zn_0_, the counts in NF+Zn_low_ collapse to levels similar to control. NF+Zn_high_ has lower counts than its fermented counterpart. Members of the genus *Spiroplasma* on the other hand, were detected in all control larvae, but mostly absent from larvae fed spinach (apart from 3). Moreover, members of the genus *Enterobacter* were present in almost all larvae fed non-fermented spinach, but were not detected in the control, and mainly absent from larvae fed fermented spinach. Finally, members of the *Weissella* genus were not detected in the control but present in most larvae fed fermented spinach (where Zn_high_ seemed to increase their counts) and absent from most larvae fed non-fermented spinach (apart from 2).

The Shannon Index (alpha-diversity) was estimated to depict species diversity and a comparison can be found in Fig. S18. The Shannon index was affected by neither the interaction between the fermentation status and the zinc concentration in the diet (F_6,56_ = 1.1433, p = 0.3498), nor by the effects of fermentation status or zinc concentration as simple effects (diet: F_6,60_ = 0.3074, p = 0.7366; zinc: F_6,58_ = 0.1028, p = 0.9025). Species diversity was generally found to be low (Shannon index ≤ 2).

An analysis of the Bray-Curtis dissimilarity matrix (beta-diversity) of all the samples showed a separation by fermentation status of the spinach in the diets, as well as zinc concentration (Fig. S19 and Table S5). Namely, larvae fed control diets are separated from larvae fed each of the treatments, regardless of fermentation status or zinc fortification. Within larvae fed a non-fermented diet, there was no separation according to zinc concentration. This was however not the case in larvae fed fermented spinach, where while larvae fed no zinc fortification and Zn_low_ were not separated, larvae fed Zn_high_ were distinct from both. Larvae fed spinach absent of zinc fortification were not separated, whether the spinach in their diets was fermented or not. This was however not the case in larvae fed Zn_low_ and Zn_high_.

## Discussion

The work presented here was motivated by a growing interest in alternative sustainable sources of protein and necessity for optimizing the diets of edible insects to improve their nutritional and safety profile. Producing insects for food and feed with optimised nutritional and safety profiles can provide even more traction to an already growing field, with a projected market growth of around 8Lbillion USD by 2030 (Omuse et al., 2024). One of many advantages of insect rearing is their ability to be bred on low-cost organic side-streams and wastes (Varelas, 2019). In fact, upcycling locally produced side-streams and by-products, and their bioconversion to high-value animal protein through insect rearing is fully aligned with circular economy practices that are currently strategically favoured and promoted in the EU (EU Commission, 2014; 2020). Several studies have reported the inclusion of agricultural side-streams and by-products into insect rearing diets, showing that when nutritionally crafted, these diets are well suited to feed insects without compromising health (Gourgouta et al., 2022; Klüber et al., 2022; Kotsou et al., 2023; Kotsou et al., 2024; Noyens et al., 2023; Rumbos et al., 2022).

Fermentation of substrate ingredients has not only been used to prolong their shelf life or improve digestibility, but more recently as a platform to introduce probiotic bacteria into insect feed (Klüber et al., 2022; Kuttiyatveetil et al., 2019; Taghikhani et al., 2023; Van Campenhout, 2021). The effects of the administration of probiotics to TM larvae have received recent interest, but are mostly still limited to the addition of probiotic cultures to the diet (Häbermann *et al*., 2025; Lecocq *et al*., 2021; Rizou *et al*., 2022; Zhong *et al*., 2017). In this study diets supplemented with spontaneously fermented spinach and essential trace element zinc were implemented, showing potential for improving the microbiome of TM larvae, by introducing probiotic bacteria without compromising survival. Longer spinach fermentations (48-96h) produced a microbiota richer in potentially probiotic bacteria in fermented spinach samples (Fig. 1). This was evidenced by the development of genera of LAB, *Latilactobacillus*, *Levilactobacillus*, *Lactococcus* (mostly only in 48h fermentations) and *Enterococcus* (Nowak et al., 2021). LAB can be found in decomposing plant materials, fermented food, sourdough, and as part of the microbiome of animals, including insects (Mokoena, 2017). Various LAB species occur in the respiratory, intestinal, and genital tracts of animals (George et al., 2018). LAB can inhibit the expansion of pathogens in the gut by competing for the same nutrients (Iorizzo et al., 2020) and the products of their metabolism (e.g., organic acids, carbon dioxide, ethanol, hydrogen peroxide) can also contribute to deter pathogens (Serna-Cock et al., 2019). LAB can also produce bacteriocins, proteinaceous molecules that disturb bacterial growth (Alvarez-Sieiro et al., 2016). Gut microbiota are significantly involved in the immunomodulation of the host, and gut LAB participate in these interactions, enhancing the ratio between anti-inflammatory and proinflammatory cytokines (Foligne et al., 2007). Given this evidence it is encouraging to observe that spontaneous spinach fermentation can lead to development of LAB populations.

Single colonies isolated from fermented spinach were identified to be of *Enterococcus* (Fig. S2), of which a few bacterial species have been investigated and used as probiotics (Hanchi et al., 2018; Krawczyk et al., 2021; Krishna et al., 2022). Interestingly, Rizou and collaborators (Rizou et al., 2022) observed that a diet supplemented with *E. faecalis* produced TM larvae with improved weight and length gain, shorter time to pupation and increased protein content, while reducing crude fat and microbial load (namely, coliforms and endospores). The enrichment of *Enterococcus* in our fermented spinach may thus be of benefit to the mealworms, although we did not detect measurable impacts on growth in our study. Furthermore, longer spinach fermentations led to significant reduction of *Pseudomonas* counts in comparison to non-fermented spinach (Fig. 1 and S1). The *Pseudomonas* genus includes important plant pathogens (Xin et al., 2018), agents of food spoilage (Mellor et al., 2011) and opportunistic pathogens to animals and humans (Moradali et al., 2017). Although not every *Pseudomonas* species is pathogenic, an effective reduction of the presence of this genus could grant our spinach substrate a safer feed. It could also extend shelf life as the *Pseudomonas spp.* has been associated with spoilage of several food groups (e.g., dairy, meat, seafood, fruit and vegetables; (Bloomfield et al., 2024; Tirloni et al., 2021)).

Beta-diversity showed clear dissimilarities in microbiota composition between non-fermented and fermented spinach, promoted by the fermentation process (Fig. S3). Similarly, the microbiota of wheat bran was also bluntly different from any spinach sample.

A comprehensive nutritional and anti-nutritional evaluation of the diets was performed to assess their suitability. The experimental and control diets used had a high dry matter (DM) content (78–95.6%), while the spinach, provided to the mealworms in the respective treatments, had a low DM content (7.9-9.4%), making them a suitable source of water (Table 1). Whereas DM values are within the expected range for wheat bran-based diets (Gianotten et al., 2020; Polovinski-Horvatović, 2025; Rumbos et al., 2020; Soetemans et al., 2020), protein content was generally higher than that usually found in wheat bran alone, explained by the inclusion of soy and yeast in the base diet (Table 1; (Gianotten et al., 2020; Polovinski-Horvatović, 2025; Rumbos et al., 2020; Soetemans et al., 2020)).

Some organic acids present in spinach can have anti-nutritional effects, which could impact larval growth. Phytic acid can form strong chelates with cations (e.g., zinc, iron, calcium, magnesium), granting them unavailable for intestinal absorption, particularly in hosts with low phytase activity. Lee and colleagues (Lee et al., 2023) quantified phytic acid and oxalates of an array of green leafy greens, including spinach. As their results are based on fresh mass, our results are in the same value range for dry mass. Interestingly, having also assessed the phytic acid content of all completed diets used in the feeding trial, no phytic acid was detected. This was surprising since wheat bran, the major component of our diets, is known to have substantial amounts of this organic acid (Guo et al., 2015), suggesting that either it was already broken down or this particular wheat variety is very poor in phytic acid.

The aforementioned study (Lee et al., 2023) also determined the oxalic acid content of spinach, which was substantially higher than those found at any of the fermentation periods, especially considering that our values portray oxalic acid content in dry mass. These findings are particularly relevant as oxalic acid has strong chelating abilities and has shown to exert a negative effect on dietary mineral absorption (Noonan and Savage, 1999). Absorption of excessive soluble oxalate can lead to hyperoxaluria (i.e., high urinary oxalate excretion), which is a significant risk factor for kidney stone formation (Bargagli et al., 2020).

LAB, including those found in fermented spinach (Fig. 1 and S1), metabolize sugar and other monosaccharides to lactic acid as part of the fermentation (Wang et al., 2021), which explains the increase in lactic acid content. Furthermore, some bacterial species (including LAB) can metabolize lactic acid and use it as an additional source of energy, especially when other sugar sources are no longer available (Jiang et al., 2014), which could explain the later decline in the number of counts for these genera at 96 h of fermentation

Levulinic acid is a carbon source that is readily obtainable from lignocellulosic biomass (such as spinach) and can serve as the sole carbon source for some bacteria (e.g., *Pseudomonas*, *Stenotrophomonas*; (Rand et al., 2017)). In our study, levulinic acid was quickly metabolized, being no longer detectable at 48h, which is consistent with the degradation rate found by Habe et al. (Habe et al., 2015) for several bacterial strains.

In fermentation dominated by lactic acid bacteria the degradation of citric acid is rather the exception, strongly depending on the specific conditions of the fermentation and the microorganisms involved. Nonetheless, it can be fermented by a limited number of LAB genera (e.g., *Leuconostoc*, *Lactococcus*, *Enterococcus*, *Lactobacillus*), some of which found on the 16S sequencing of fermented spinach (Fig. 1; (Hugenholtz, 1993)). Moreover, LAB can break down malic acid through a process called malolactic fermentation, converting it into lactic acid and carbon dioxide (Kunkee, 1991). A significant decrease in malic content in spinach was observed with the start of the fermentative process, which could be explained by malolactic fermentation, also contributing to a significant increase in lactic acid.

The FAN content can serve as an indicator of fermentation activity, as low values can indicate a lack of nutrients, which can lead to incomplete or slow fermentations (Schaan and Hughes, 2024). Data shows that FAN values doubled after 48h fermentations when compared to non-fermented, falling back to the initial value after 96h (Table 3). This pattern is common, where FAN peaks early, as proteolytic microorganisms (e.g., *Lactobacillus*, *Leuconostoc*) are active, then falls as microbial populations grow and assimilate nitrogen (Eppendorfer and Bille, 1996). This is consistent with the amino acid profile assessment of non-fermented and fermented spinach (Table 4), where most amino acids (especially glutamic acid, proline, glycine and valine) increase dramatically at 24-48h, due to microbial proteolysis. This was, again, followed by a sharp decrease at 96h, as they are consumed as energy/nitrogen sources or degraded into other metabolites by fermenting microorganisms.

Volatile compound analysis strongly suggests a healthy fermentative process, with presence of LAB populations, further supported by the absence of volatile compounds in non-fermented spinach (Table 5), indicating that they are products of fermentation rather than pre-existing in the spinach. At the early stage of fermentation (i.e., 24h) several compounds emerge: ethyl acetate, 2-methylbutanal, 3-methylbutanal, hexanal, 2-butanone. Aldehydes 2-methylbutanal and 3-methylbutanal are naturally found in spinach, indicating active amino acid metabolism (of isoleucine and leucine, respectively) by bacteria, such as LAB (Brandsma et al., 2024; Lee et al., 2016; Masanetz et al., 1998), increasing throughout the length of the fermentation. Interestingly, fermentation of oats and soybeans with LAB species led to the production of ethyl acetate, through the reaction of ethanol with acetic acid (Lee et al., 2016; Li et al., 2024), which was also observed in our samples, peaking at 48h, but completely consumed, or volatised, by 96h. Hexanal is an aldehyde, shown to be present in spinach (especially when boiled), produced from the oxidation of fatty acids (Masanetz et al., 1998). Ketones, like 2-butanone, are commonly found in LAB fermented products, having a major influence on aroma (Gallegos et al., 2017).

The later stage of the fermentation (i.e., 96h) was highlighted by significant increase of 3-methyl-1-butanol and benzaldehyde, as well as emergence of dimethyl disulfide. All these 3 compounds can be synthesized from the metabolism of branched-chain amino acids present in organic materials by LAB. Methyl alcohols, such as 3-methyl-1-butanol, from catabolism of leucine and isoleucine (Chen et al., 2019; Schoondermark-Stolk et al., 2006), benzaldehyde using phenylalanine as a metabolite (Kunjapur and Prather, 2015; Nierop Groot and de Bont, 1998) and dimethyl disulfide from the degradation of methionine (Lee et al., 2016). This correlates with the significant reduction in the content of those amino acids at 96h (Table 4).

The experimental conditions depicted in this study had negligible impact on survival (Fig. 2B), however spinach-based diets were observed to lead to a slower larval growth rate, regardless of fermentation, when compared to control (Fig. 2A and S13). Such can be partially explained by contrasting nutritional differences between spinach and the apple used as control, the latter boasting substantially higher calorie and carbohydrate count (USDA, 2025a; b). Having access to a higher supply of readily available energy could therefore improve growth rate. It is also important to notice that the feeding trial was terminated once the first pupae were observed, which were found in the control group. Given that all the treatments registered negligible effects on survival, it is fair to assume that if allowed to achieve full larval maturity (i.e., reach pupae phase), the experimental groups would have gained similar weight to that of the control.

While supplementation with low dosages of zinc did not result in significant weight gain differences to that of control, rearing with Zn_high_ diets led to significantly lower weight gain (Fig. 2A and S13). Thus, it seems likely that dietary efficiency is closely related to micronutrient composition, as developmental retardation in zinc feeding trials has been previously observed for terrestrial insects, including TM (Gintenreiter et al., 1993; Keil et al., 2020; Noret et al., 2007). This could be the result of a systemic intoxication promoted by elevated zinc feeding (Nishito and Kambe, 2018; Sandstrom, 2001). Nonetheless, TM like any other organism, requires a certain amount of supplemental zinc, previously determined by Fraenkel (Fraenkel, 1958) to be 6 ppm for optimal larval growth, which was already exceeded in control diets.

Consistent with previous findings (Bednarska and Swiatek, 2016; Keil et al., 2020), we observed a clear dosage-dependent accumulation of zinc in mealworms based on the amount of zinc added to the diet (Fig. 3B and S15). BAF decreased with higher dietary zinc, notably at the Zn_high_ diet, indicating reduced accumulation and suggesting internal zinc regulation (Fig. 3C and S16). TM appears capable of adjusting zinc levels through homeostatic mechanisms (Dallinger, 1993; Diener et al., 2015; Keil et al., 2020). This adaptation could be the reason why, despite retarding growth, a higher zinc dosage did not increase mortality, limiting its toxicity.

Differences in the nutritional and bacterial composition of the different diets translated into differences in the bacterial communities found in TM larvae (Fig. 4). Whereas the experimental control heavily featured *Spiroplasma* counts, these are mostly absent from spinach-treated groups (Fig. 4 and S17). *Spiroplasma* species are considered opportunistic symbionts in multiple insect species and abundantly found in TM microbiota (Jung et al., 2014; Mamtimin et al., 2023; Montalban et al., 2022). By keeping a symbiotic equilibrium, *Spiroplasma* is usually not harmful in the mealworm gut (Jung et al., 2014). The absence of *Spiroplasma* from most spinach-treated groups seems to indicate that diet is the reason for this outcome, which could be promoting competitive microbiota or an unfavourable nutrient profile.

In accordance with previous studies *Staphylococcus* species can be found in the microbiota of TM fed bran-based diets (Lou et al., 2021; Mamtimin et al., 2023; Montalban et al., 2022), sometimes in great abundance (Savio et al., 2024). The larvae in our experiment harboured bacteria of the *Staphylococcus* genus, which, although present in all experimental groups, saw its counts significantly increased in spinach-supplemented diets, regardless of fermentation, compared to control. Although *Staphylococcus* species have notoriously been associated with pathogenic infections in multiple taxa, including insects and mammals (Andrade-Oliveira et al., 2023; Clement et al., 2005; McGonigle et al., 2016), interestingly, some have also been shown to be probiotic in *Bombyx mori* (Saranya et al., 2019).

Larvae fed fermented spinach presented increased counts of LAB, specifically *Lactococcus* and *Weissella*, genera notoriously associated with probiotic species (Franz and Holzapfel, 2011). Species from these genera can increase the presence of antimicrobial molecules in the gut, such as bacteriocins, organic acids, hydrogen peroxide, acetaldehyde, acetoin, and carbon dioxide (Arena et al., 2018), which may confer protection to insects from entomopathogen infections (Savio et al., 2024). A prime example is Nisin, bacteriocin produced principally by *Lactococcus lactis* (O’Connor et al., 2015), which has been commercially available for over 60 years (Cotter et al., 2005) and is mainly active against Gram-positive bacteria (e.g., *Listeria* and *Staphylococcus*), and the spore forming bacteria (e.g., *Bacillus* and *Clostridium*; (And and Hoover, 2003)).

Weissella was the only genus whose counts were affected by zinc supplementation in interaction with the fermentation status of the spinach in the diet. In the presence of fermented spinach, the highest zinc dosage led to increased *Weissella* counts. The effects of zinc on insect gut microbiome have been insufficiently explored, focusing on other species (namely piglets), where data indicates that the effects tend to be species and dosage-specific (Skalny et al., 2021). Interestingly, Vahjen et al. (Vahjen et al., 2011) found high dosages of zinc oxide to modulate the enterobacterial composition of piglets, significantly increasing the abundancy of *Weissella* species, *cibaria* and *confusa*, which have more recently been described for their probiotic potential (Quattrini et al., 2020).

Although there were no significant differences in alpha diversity among treatments (Fig. S18), there were differences in beta diversity (Fig. S19 and Table S5). The microbiota composition of the control group is significantly different from all treatment groups, regardless of fermentation or zinc supplementation. This is also the case for the larvae fed the fermented spinach containing diet with high levels of zinc. Interestingly, none of the genera found in the larvae were detected in the diet components, suggesting that the effect of the fermentation status of the diet on TM microbiota is indirect. This clearly illustrates a priming effect on microbiota introduced both by spinach feeding and zinc supplementation, at both lower and higher dosages, leading to the proliferation of probiotic-associated genera. Thus, here there seems to be a multifactorial requirement for microbiota composition change, encompassing spinach, fermentation and zinc.

## Conclusions

This work promoted and identified beneficial microbiota changes in spinach originated from spontaneous fermentation, which do not require a starter colony. A healthy fermentation process was illustrated with a comprehensive nutritional study encompassing several elements, where no evidence of limiting anti-nutritional factors was reported. As a result, significantly increased counts of LAB genera were found in longer fermentations, as well as repression of potentially hazardous *Pseudomonas* genus.

A 35-day feeding trial using diets supplemented with fermented spinach and zinc was performed, with negligible effects on TM larvae survival. Despite spinach and higher dosages of zinc leading to reduced growth rates, potentially due caloric deficiencies and some level of toxicity, respectively, the experimental diets led to significant changes in microbiota of larvae, when compared to control. Fermented spinach-treated groups revealed significantly increased counts of LAB genera, namely *Lactococcus* and *Weisella*, further promoted by higher zinc concentration for the latter. The impact of spinach-treated diets, especially when fermented, and zinc supplementation was also evident in microbiota diversity, promoting significant divergencies when compared to control. Microbiota diversity on its own is not inherently positive (Häbermann *et al*., 2025) but given that we have found evidence of increased probiotic populations resulting from spinach-fermented diets, especially when fortified with Zn_high_, these findings seem encouraging.

## Conflicts of interest

The authors declare no conflicts of interest.

## Funding

This work was funded by an IBB ProValid grant (VAL128/2023) to CR. ICP-MS was supported by a grant from the Foundation for Food Safety and Consumer Protection (Stiftung für Lebensmittelsicherheit und Verbraucherschutz) to HH.

## Notes

### Competing Interest Statement

The authors have declared no competing interest.

