## Supplementary information for "Fermentation of Agricultural By-products and Zinc Supplementation: A Synergistic Approach to Mealworm Microbiome Optimization"

### Slide 1
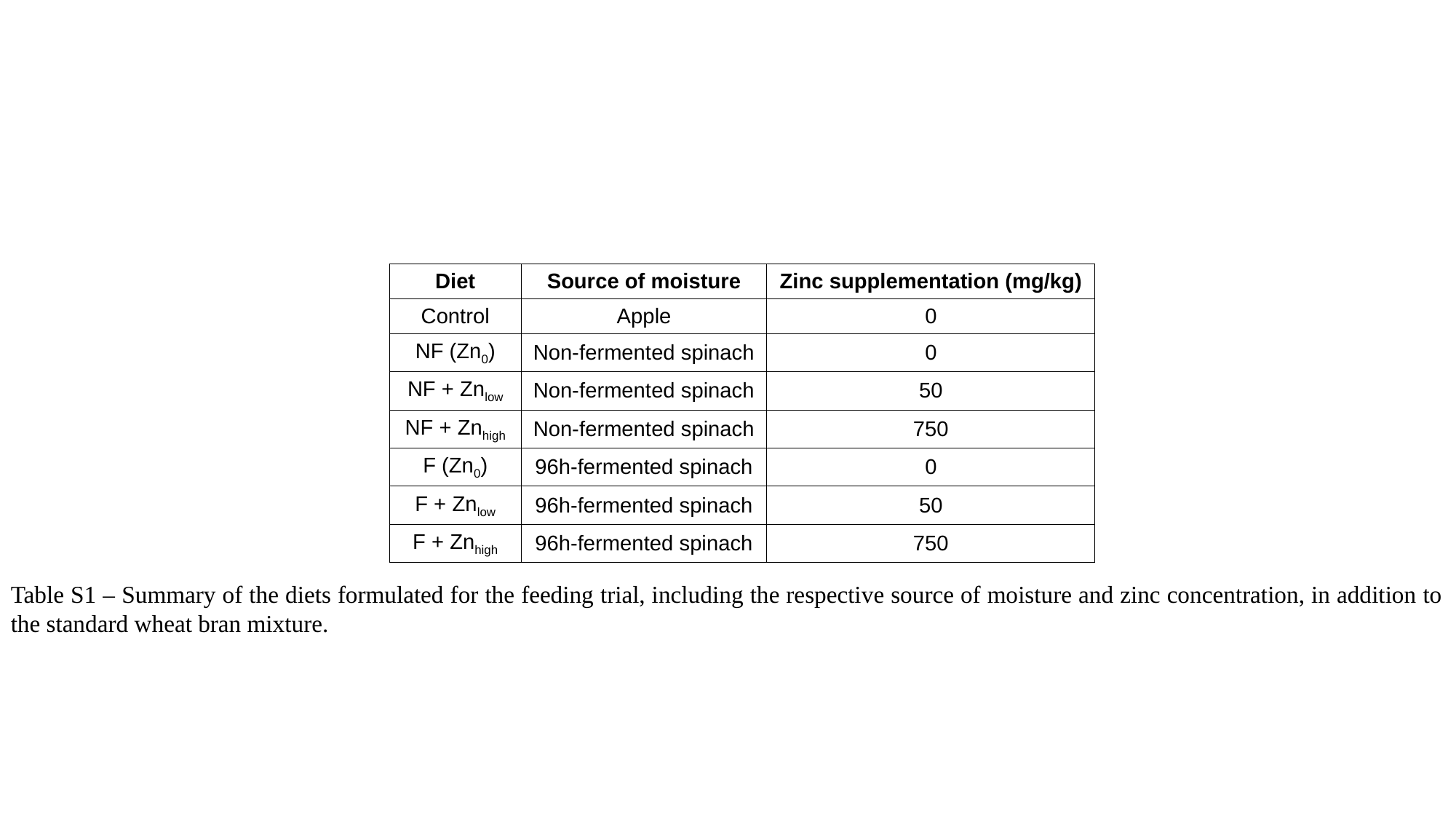

| Diet | Source of moisture | Zinc supplementation (mg/kg) |
| --- | --- | --- |
| Control | Apple | 0 |
| NF (Zn_0_) | Non-fermented spinach | 0 |
| NF + Zn_low_ | Non-fermented spinach | 50 |
| NF + Zn_high_ | Non-fermented spinach | 750 |
| F (Zn_0_) | 96h-fermented spinach | 0 |
| F + Zn_low_ | 96h-fermented spinach | 50 |
| F + Zn_high_ | 96h-fermented spinach | 750 |
Table S1 – Summary of the diets formulated for the feeding trial, including the respective source of moisture and zinc concentration, in addition to the standard wheat bran mixture.

### Slide 2
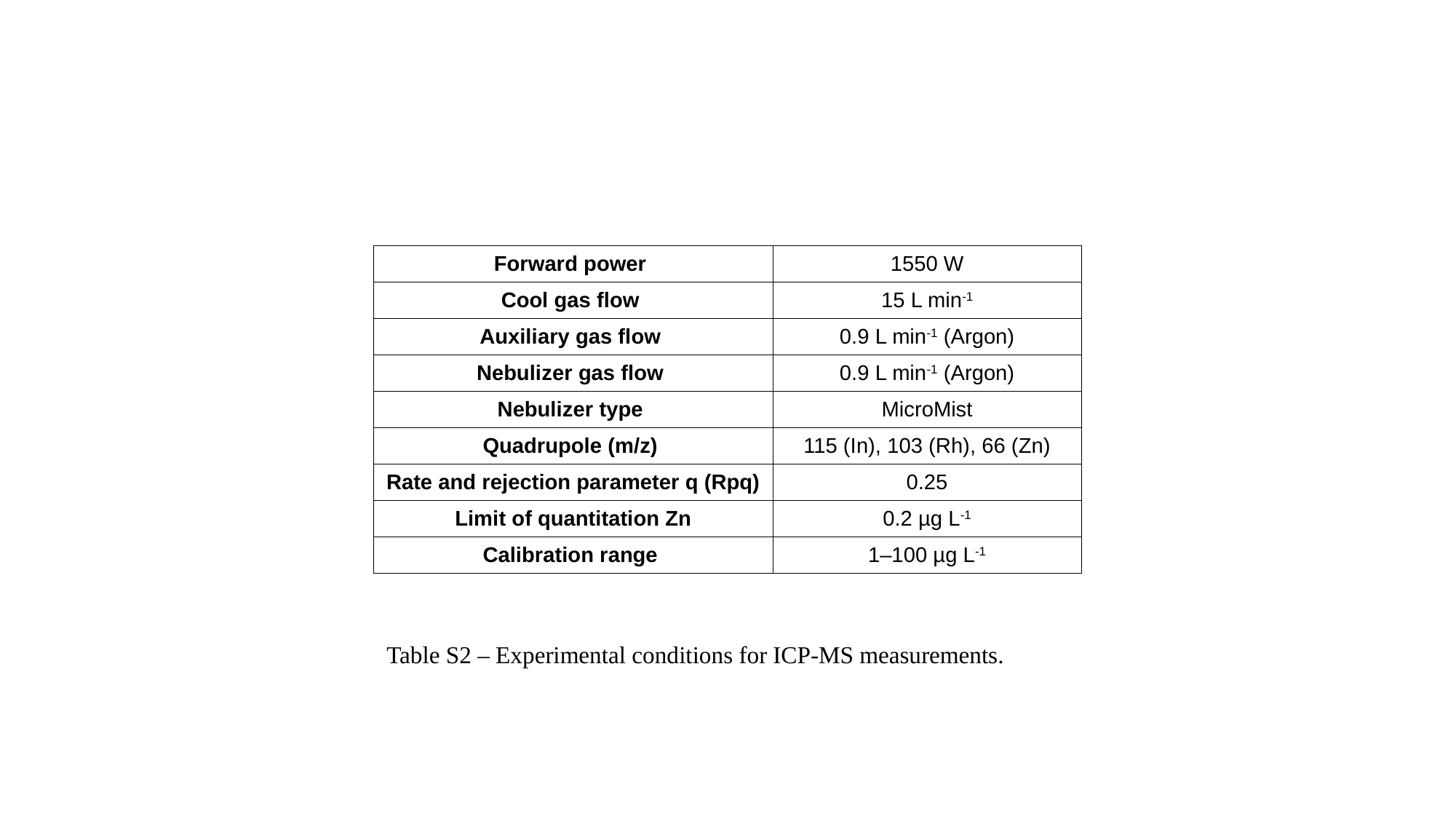

| Forward power | 1550 W |
| --- | --- |
| Cool gas flow | 15 L min^-1^ |
| Auxiliary gas flow | 0.9 L min^-1^ (Argon) |
| Nebulizer gas flow | 0.9 L min^-1^ (Argon) |
| Nebulizer type | MicroMist |
| Quadrupole (m/z) | 115 (In), 103 (Rh), 66 (Zn) |
| Rate and rejection parameter q (Rpq) | 0.25 |
| Limit of quantitation Zn | 0.2 µg L^-1^ |
| Calibration range | 1–100 µg L^-1^ |
Table S2 – Experimental conditions for ICP-MS measurements.

### Slide 3
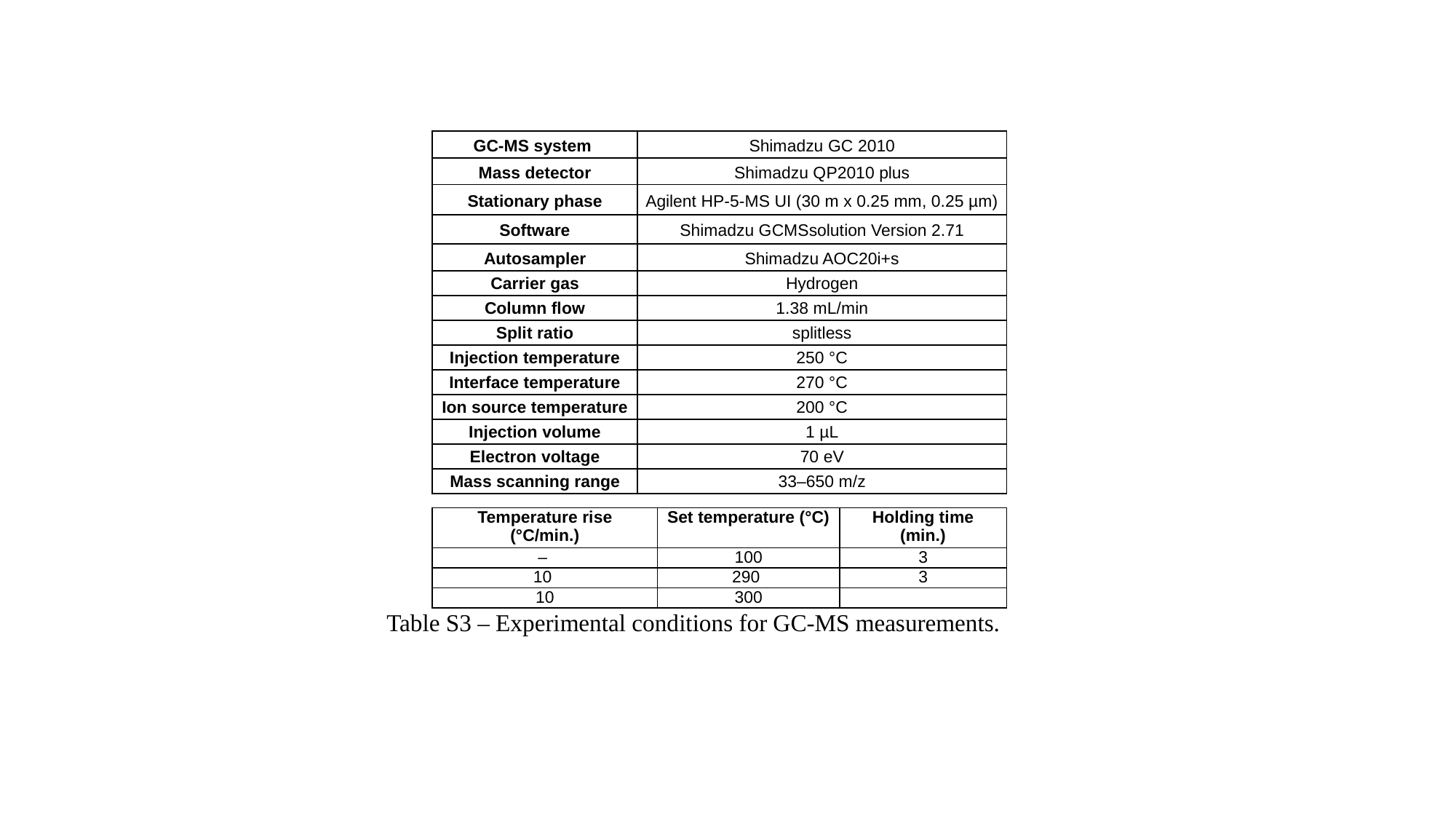

| GC-MS system | Shimadzu GC 2010 |
| --- | --- |
| Mass detector | Shimadzu QP2010 plus |
| Stationary phase | Agilent HP-5-MS UI (30 m x 0.25 mm, 0.25 µm) |
| Software | Shimadzu GCMSsolution Version 2.71 |
| Autosampler | Shimadzu AOC20i+s |
| Carrier gas | Hydrogen |
| Column flow | 1.38 mL/min |
| Split ratio | splitless |
| Injection temperature | 250 °C |
| Interface temperature | 270 °C |
| Ion source temperature | 200 °C |
| Injection volume | 1 µL |
| Electron voltage | 70 eV |
| Mass scanning range | 33–650 m/z |
| Temperature rise (°C/min.) | Set temperature (°C) | Holding time (min.) |
| --- | --- | --- |
| – | 100 | 3 |
| 10 | 290 | 3 |
| 10 | 300 | |
Table S3 – Experimental conditions for GC-MS measurements.

### Slide 4
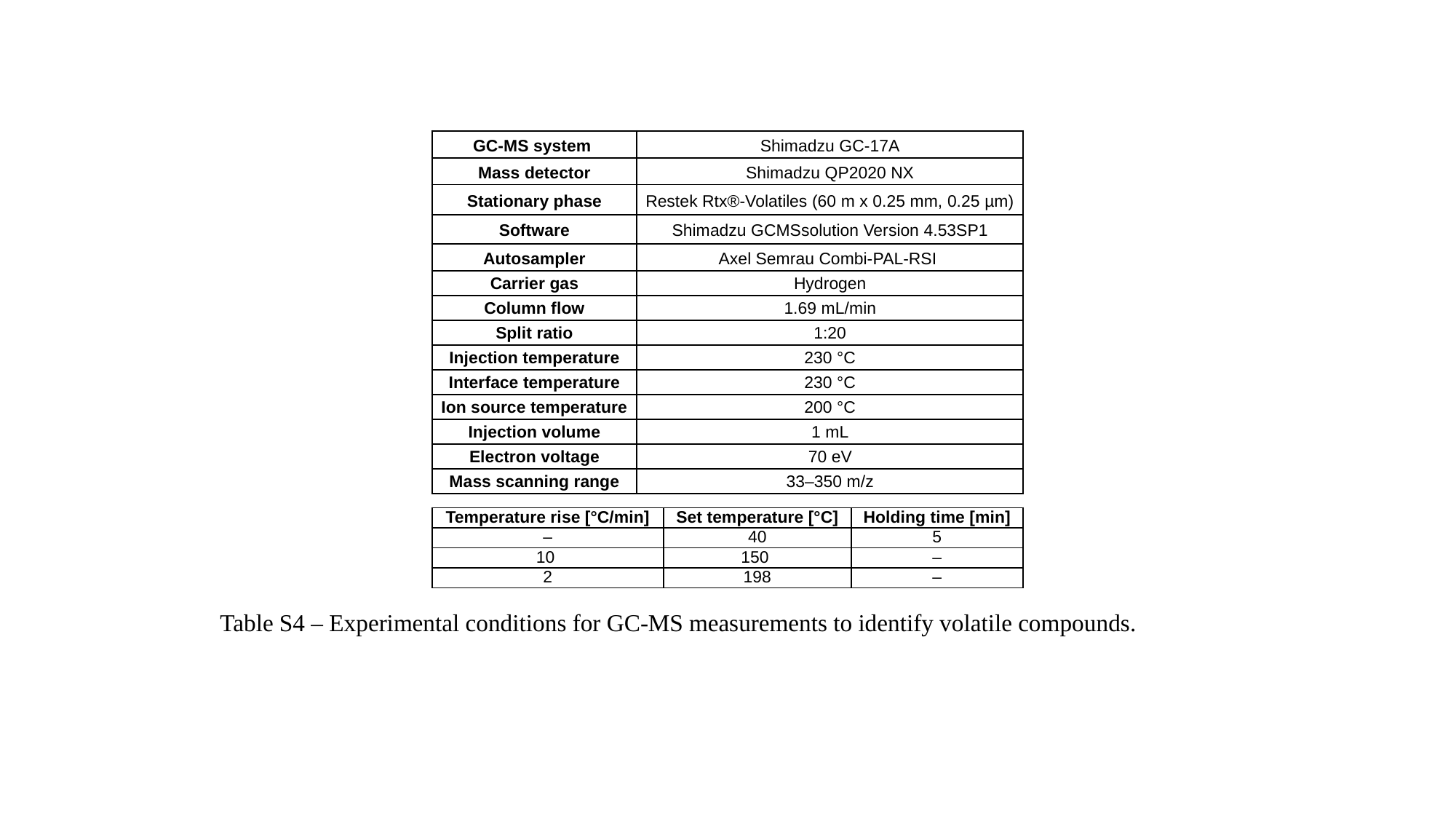

| GC-MS system | Shimadzu GC-17A |
| --- | --- |
| Mass detector | Shimadzu QP2020 NX |
| Stationary phase | Restek Rtx®-Volatiles (60 m x 0.25 mm, 0.25 µm) |
| Software | Shimadzu GCMSsolution Version 4.53SP1 |
| Autosampler | Axel Semrau Combi-PAL-RSI |
| Carrier gas | Hydrogen |
| Column flow | 1.69 mL/min |
| Split ratio | 1:20 |
| Injection temperature | 230 °C |
| Interface temperature | 230 °C |
| Ion source temperature | 200 °C |
| Injection volume | 1 mL |
| Electron voltage | 70 eV |
| Mass scanning range | 33–350 m/z |
| Temperature rise [°C/min] | Set temperature [°C] | Holding time [min] |
| --- | --- | --- |
| – | 40 | 5 |
| 10 | 150 | – |
| 2 | 198 | – |
Table S4 – Experimental conditions for GC-MS measurements to identify volatile compounds.

### Slide 5
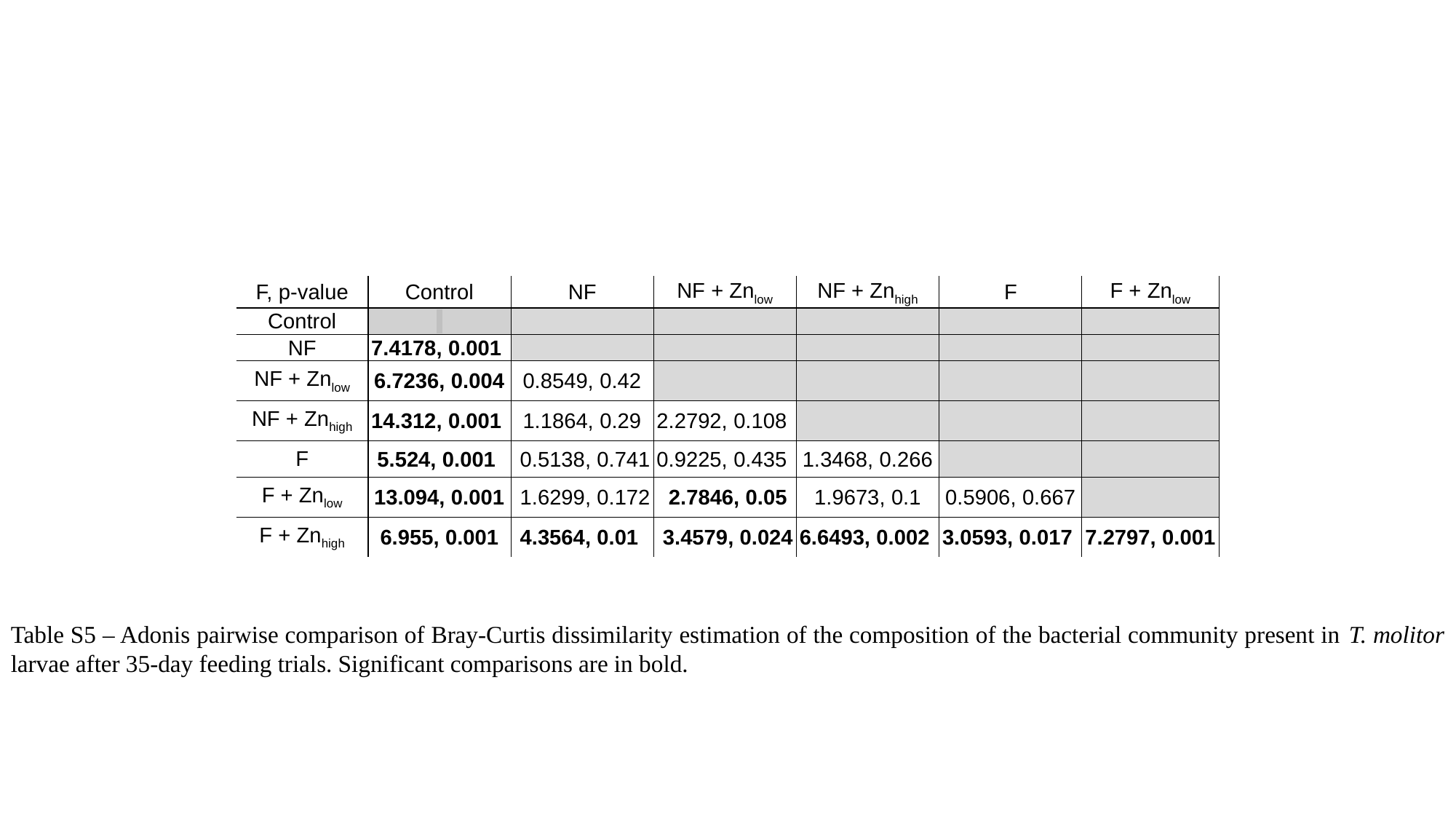

| F, p-value | Control | NF | NF + Zn_low_ | NF + Zn_high_ | F | F + Zn_low_ |
| --- | --- | --- | --- | --- | --- | --- |
| Control | | | | | | |
| NF | 7.4178, 0.001 | | | | | |
| NF + Zn_low_ | 6.7236, 0.004 | 0.8549, 0.42 | | | | |
| NF + Zn_high_ | 14.312, 0.001 | 1.1864, 0.29 | 2.2792, 0.108 | | | |
| F | 5.524, 0.001 | 0.5138, 0.741 | 0.9225, 0.435 | 1.3468, 0.266 | | |
| F + Zn_low_ | 13.094, 0.001 | 1.6299, 0.172 | 2.7846, 0.05 | 1.9673, 0.1 | 0.5906, 0.667 | |
| F + Zn_high_ | 6.955, 0.001 | 4.3564, 0.01 | 3.4579, 0.024 | 6.6493, 0.002 | 3.0593, 0.017 | 7.2797, 0.001 |
Table S5 – Adonis pairwise comparison of Bray-Curtis dissimilarity estimation of the composition of the bacterial community present in T. molitor larvae after 35-day feeding trials. Significant comparisons are in bold.

### Slide 6
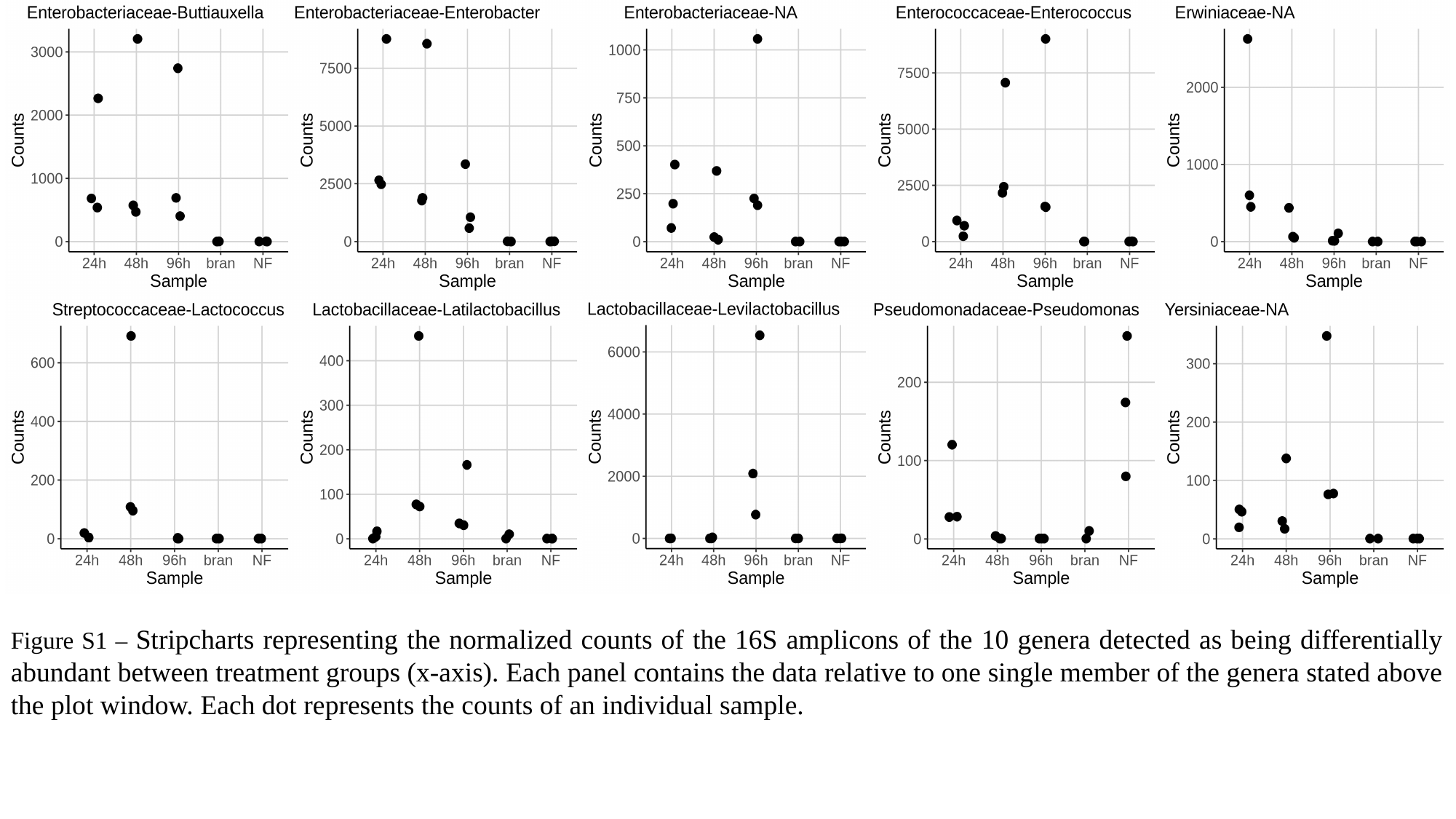

Figure S1 – Stripcharts representing the normalized counts of the 16S amplicons of the 10 genera detected as being differentially abundant between treatment groups (x-axis). Each panel contains the data relative to one single member of the genera stated above the plot window. Each dot represents the counts of an individual sample.

### Slide 7
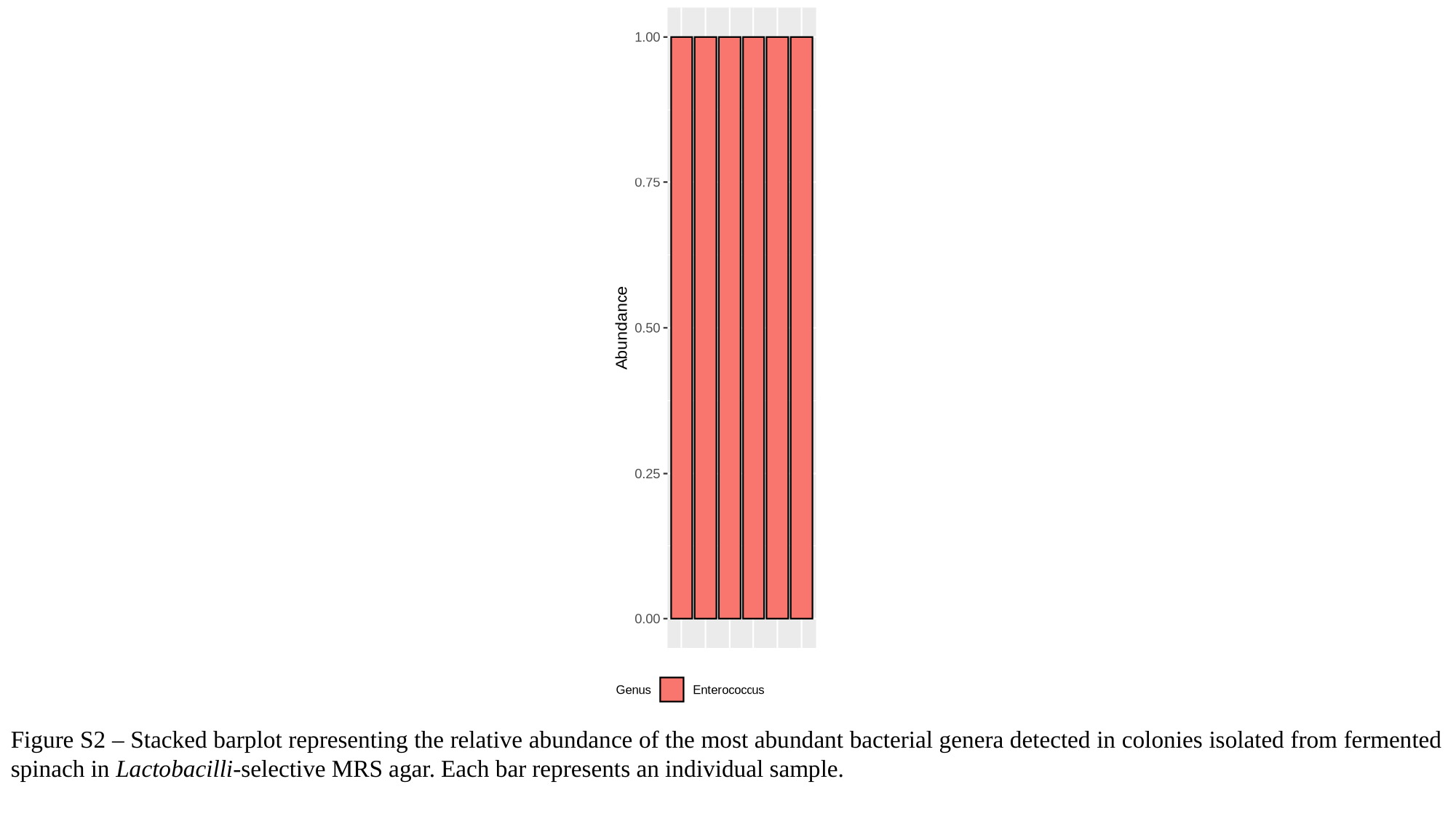

Figure S2 – Stacked barplot representing the relative abundance of the most abundant bacterial genera detected in colonies isolated from fermented spinach in Lactobacilli-selective MRS agar. Each bar represents an individual sample.

### Slide 8
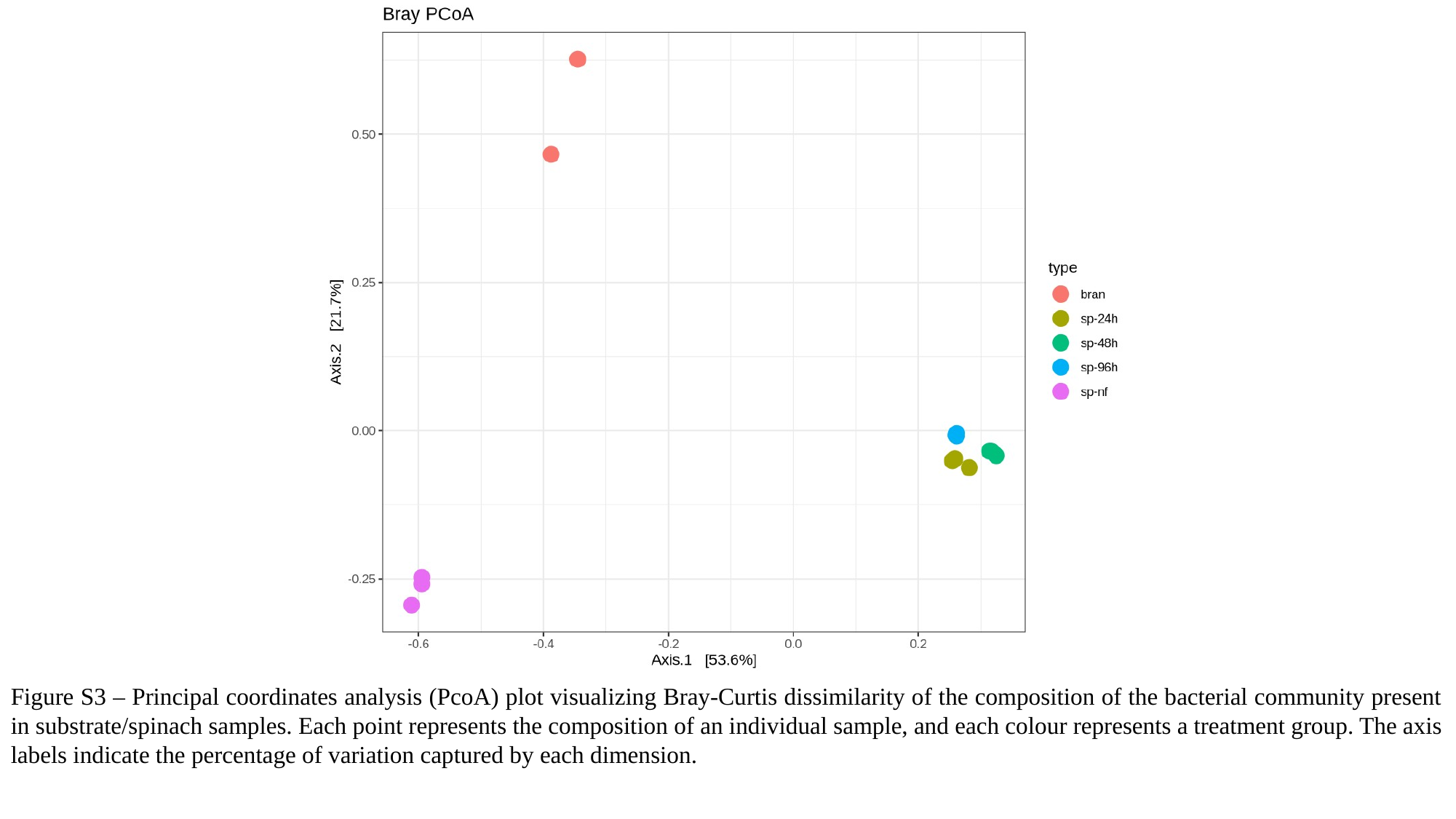

Figure S3 – Principal coordinates analysis (PcoA) plot visualizing Bray-Curtis dissimilarity of the composition of the bacterial community present in substrate/spinach samples. Each point represents the composition of an individual sample, and each colour represents a treatment group. The axis labels indicate the percentage of variation captured by each dimension.

### Slide 9
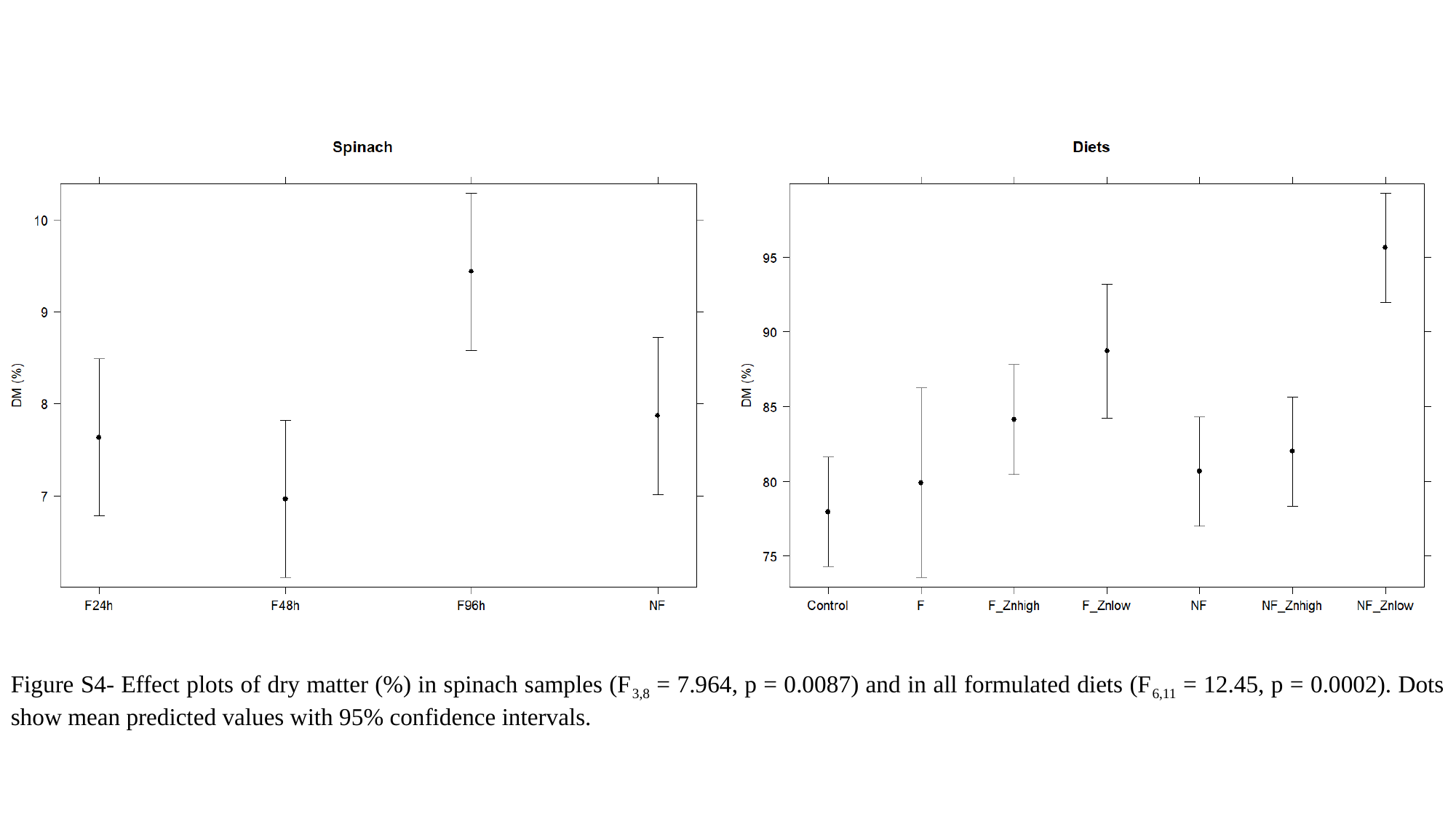

Figure S4- Effect plots of dry matter (%) in spinach samples (F_3,8_ = 7.964, p = 0.0087) and in all formulated diets (F_6,11_ = 12.45, p = 0.0002). Dots show mean predicted values with 95% confidence intervals.

### Slide 10
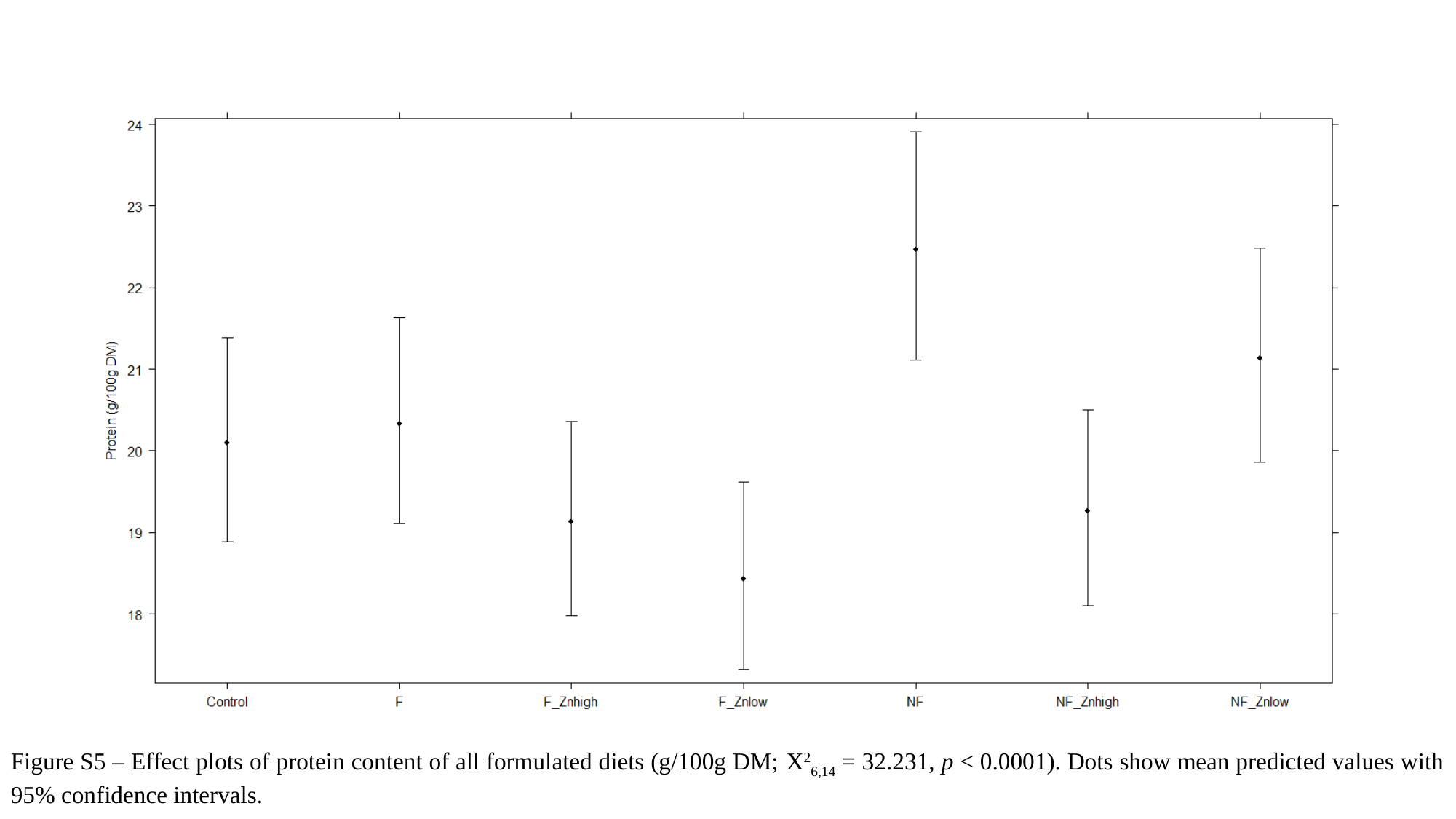

Figure S5 – Effect plots of protein content of all formulated diets (g/100g DM; X^2^_6,14_ = 32.231, p < 0.0001). Dots show mean predicted values with 95% confidence intervals.

### Slide 11
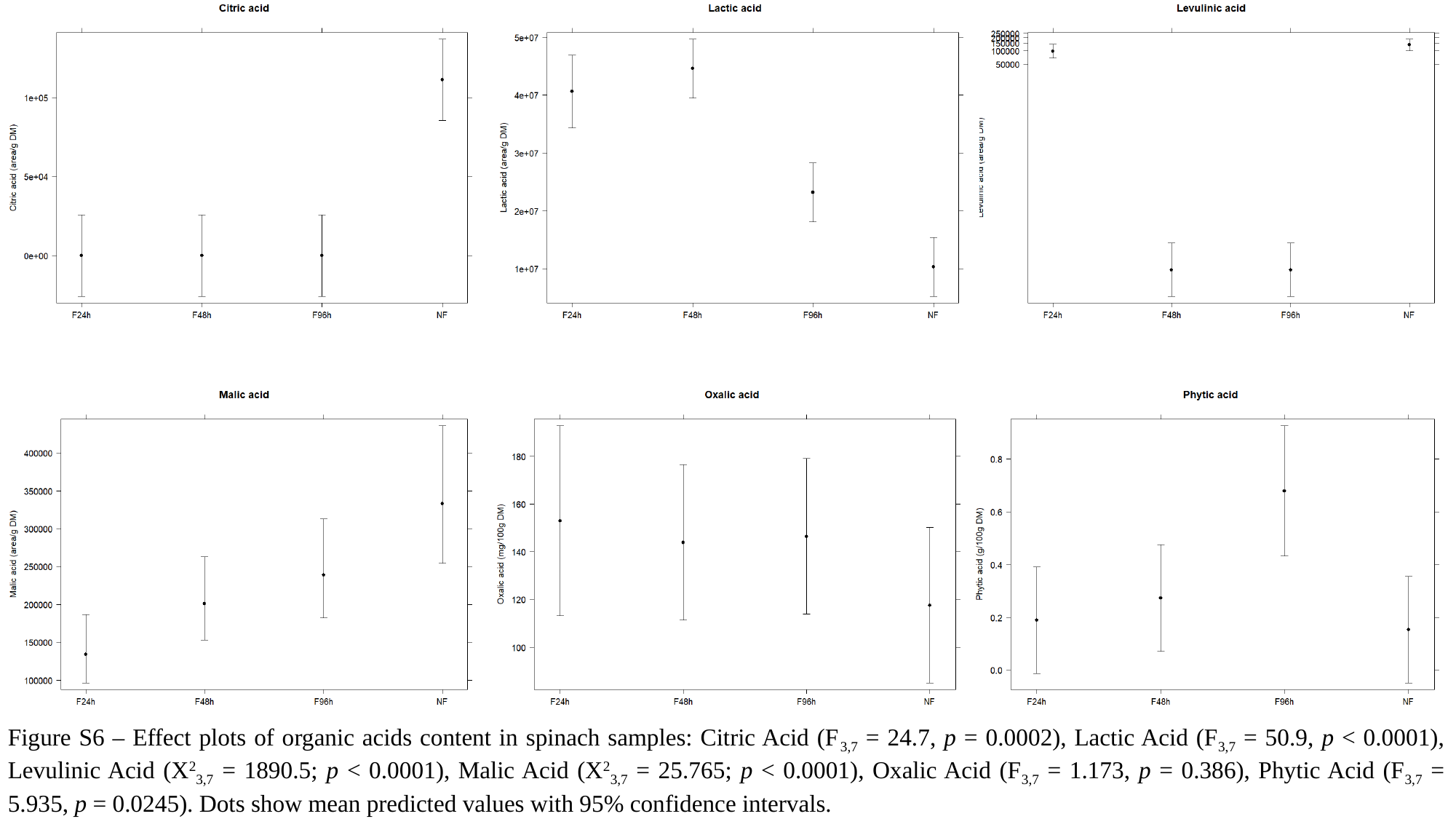

Figure S6 – Effect plots of organic acids content in spinach samples: Citric Acid (F_3,7_ = 24.7, p = 0.0002), Lactic Acid (F_3,7_ = 50.9, p < 0.0001), Levulinic Acid (X^2^_3,7_ = 1890.5; p < 0.0001), Malic Acid (X^2^_3,7_ = 25.765; p < 0.0001), Oxalic Acid (F_3,7_ = 1.173, p = 0.386), Phytic Acid (F_3,7_ = 5.935, p = 0.0245). Dots show mean predicted values with 95% confidence intervals.

### Slide 12
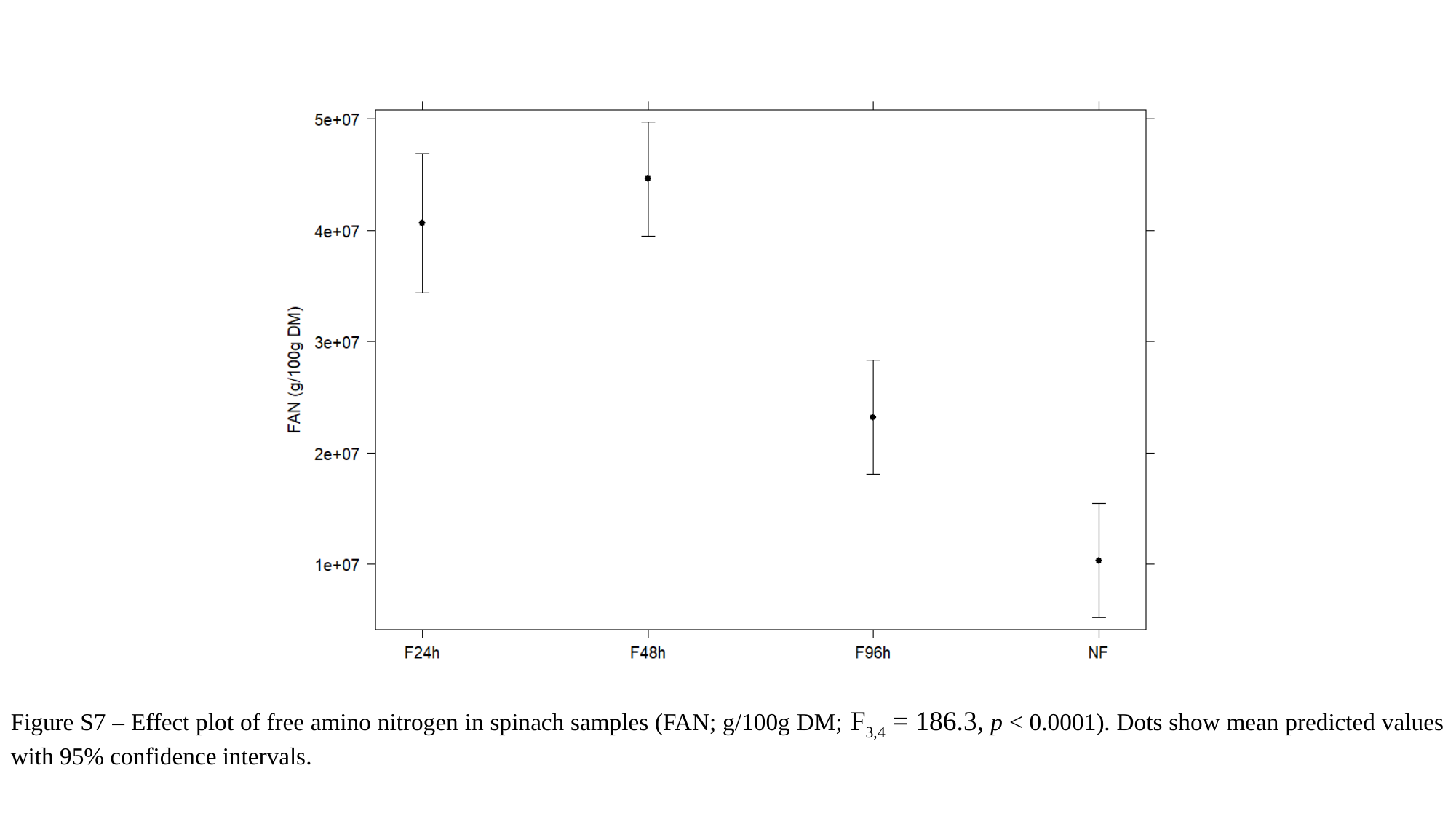

Figure S7 – Effect plot of free amino nitrogen in spinach samples (FAN; g/100g DM; F_3,4_ = 186.3, p < 0.0001). Dots show mean predicted values with 95% confidence intervals.

### Slide 13
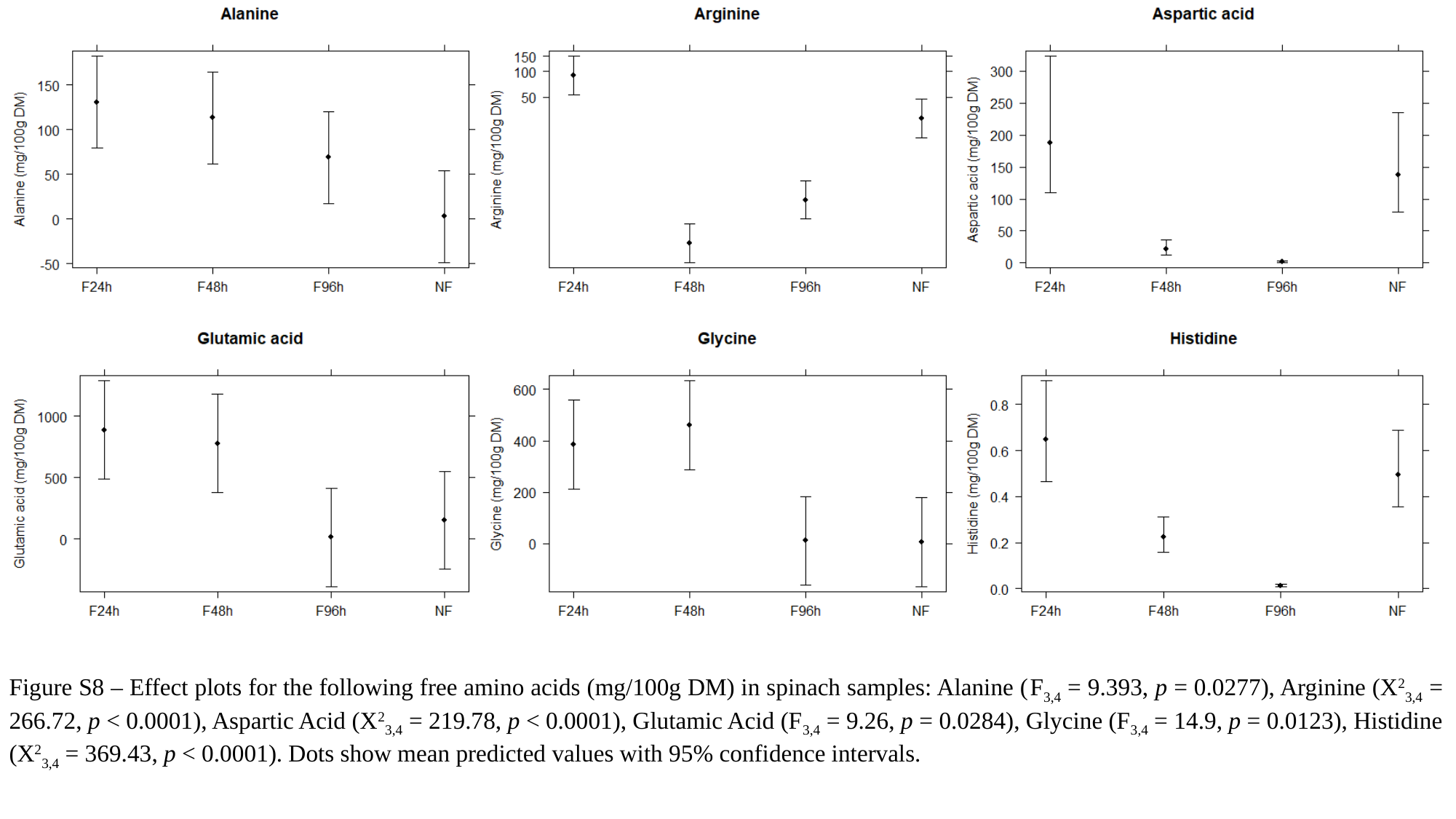

Figure S8 – Effect plots for the following free amino acids (mg/100g DM) in spinach samples: Alanine (F_3,4_ = 9.393, p = 0.0277), Arginine (X^2^_3,4_ = 266.72, p < 0.0001), Aspartic Acid (X^2^_3,4_ = 219.78, p < 0.0001), Glutamic Acid (F_3,4_ = 9.26, p = 0.0284), Glycine (F_3,4_ = 14.9, p = 0.0123), Histidine (X^2^_3,4_ = 369.43, p < 0.0001). Dots show mean predicted values with 95% confidence intervals.

### Slide 14
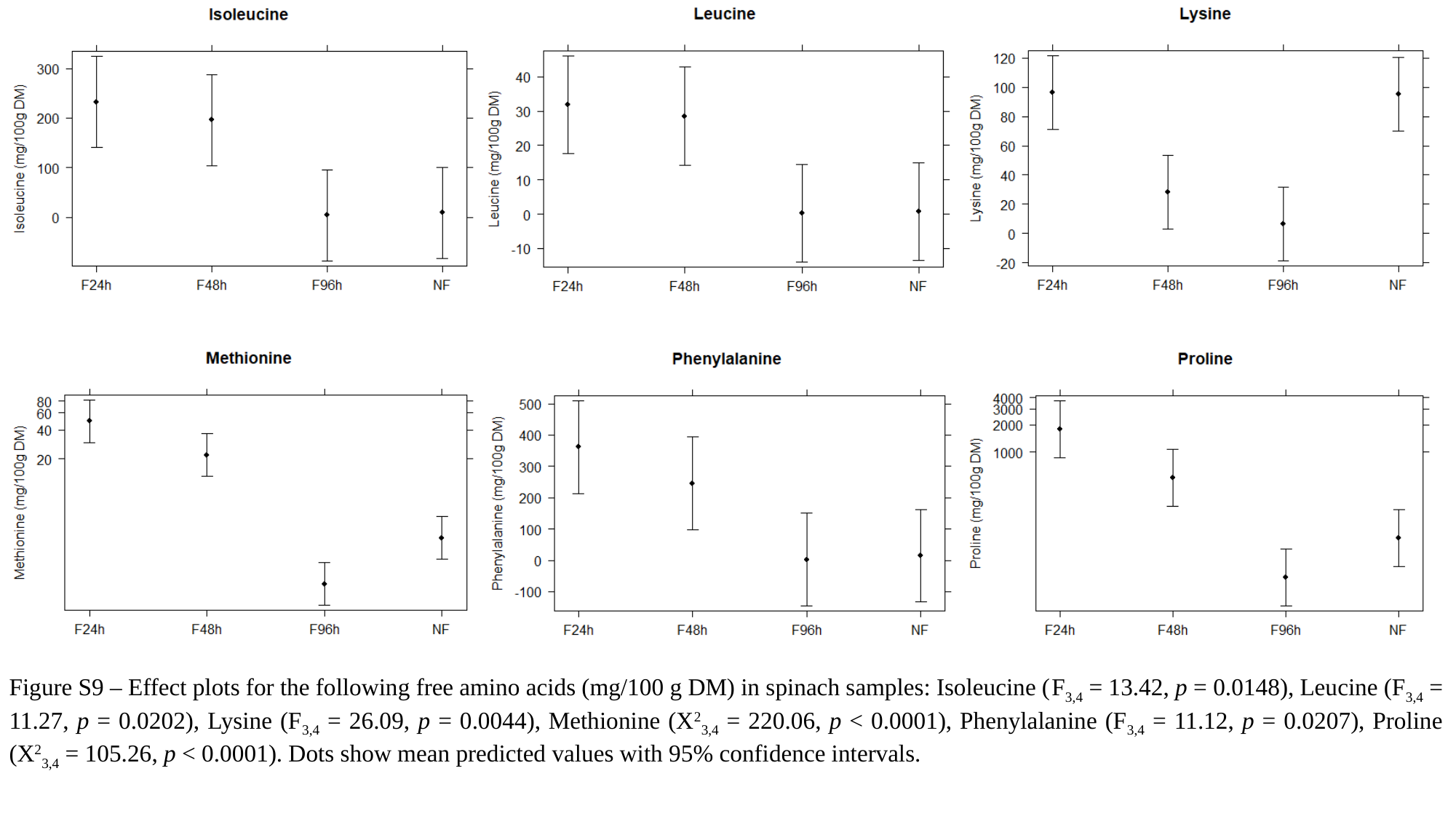

Figure S9 – Effect plots for the following free amino acids (mg/100 g DM) in spinach samples: Isoleucine (F_3,4_ = 13.42, p = 0.0148), Leucine (F_3,4_ = 11.27, p = 0.0202), Lysine (F_3,4_ = 26.09, p = 0.0044), Methionine (X^2^_3,4_ = 220.06, p < 0.0001), Phenylalanine (F_3,4_ = 11.12, p = 0.0207), Proline (X^2^_3,4_ = 105.26, p < 0.0001). Dots show mean predicted values with 95% confidence intervals.

### Slide 15
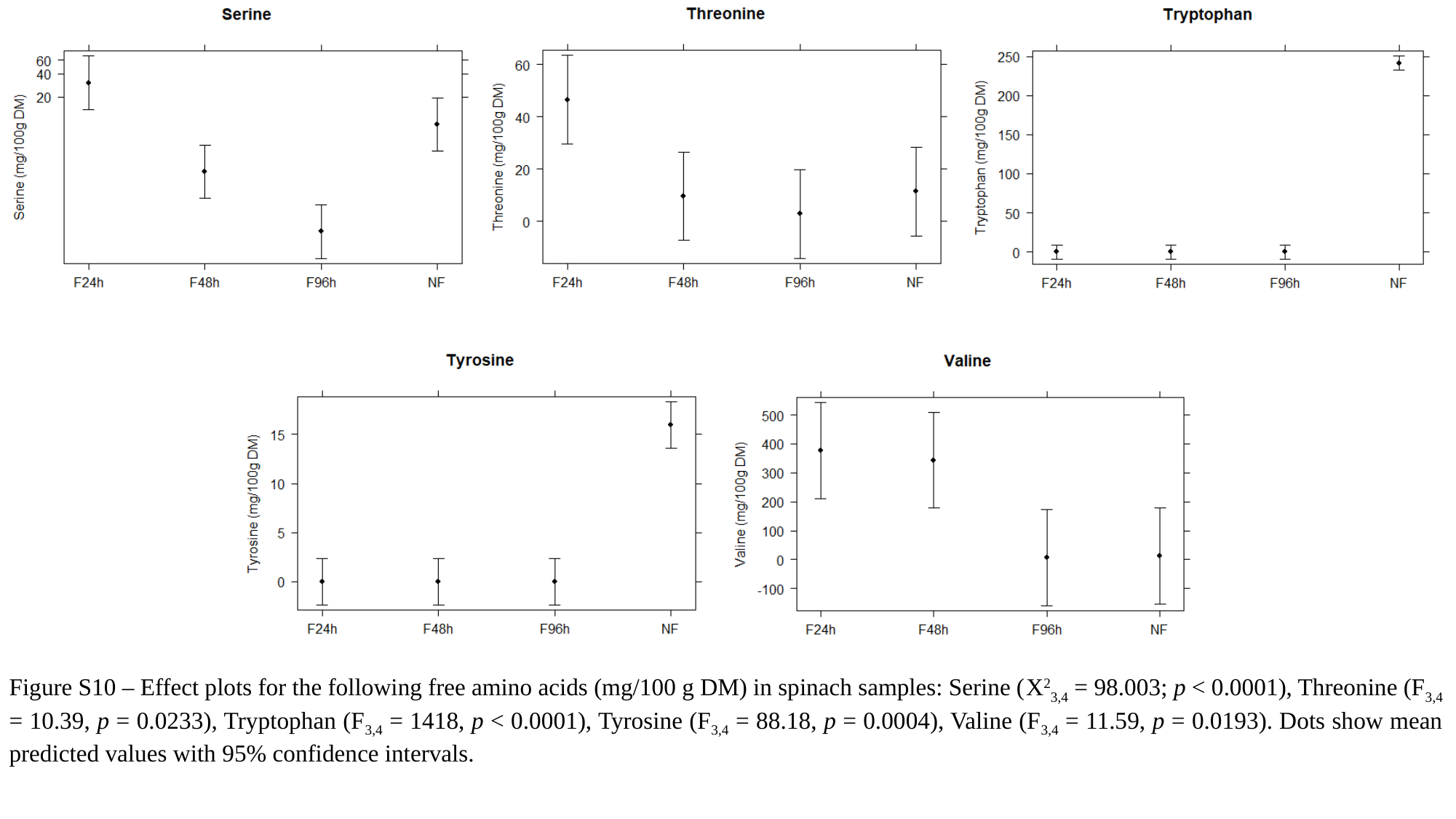

Figure S10 – Effect plots for the following free amino acids (mg/100 g DM) in spinach samples: Serine (X^2^_3,4_ = 98.003; p < 0.0001), Threonine (F_3,4_ = 10.39, p = 0.0233), Tryptophan (F_3,4_ = 1418, p < 0.0001), Tyrosine (F_3,4_ = 88.18, p = 0.0004), Valine (F_3,4_ = 11.59, p = 0.0193). Dots show mean predicted values with 95% confidence intervals.

### Slide 16
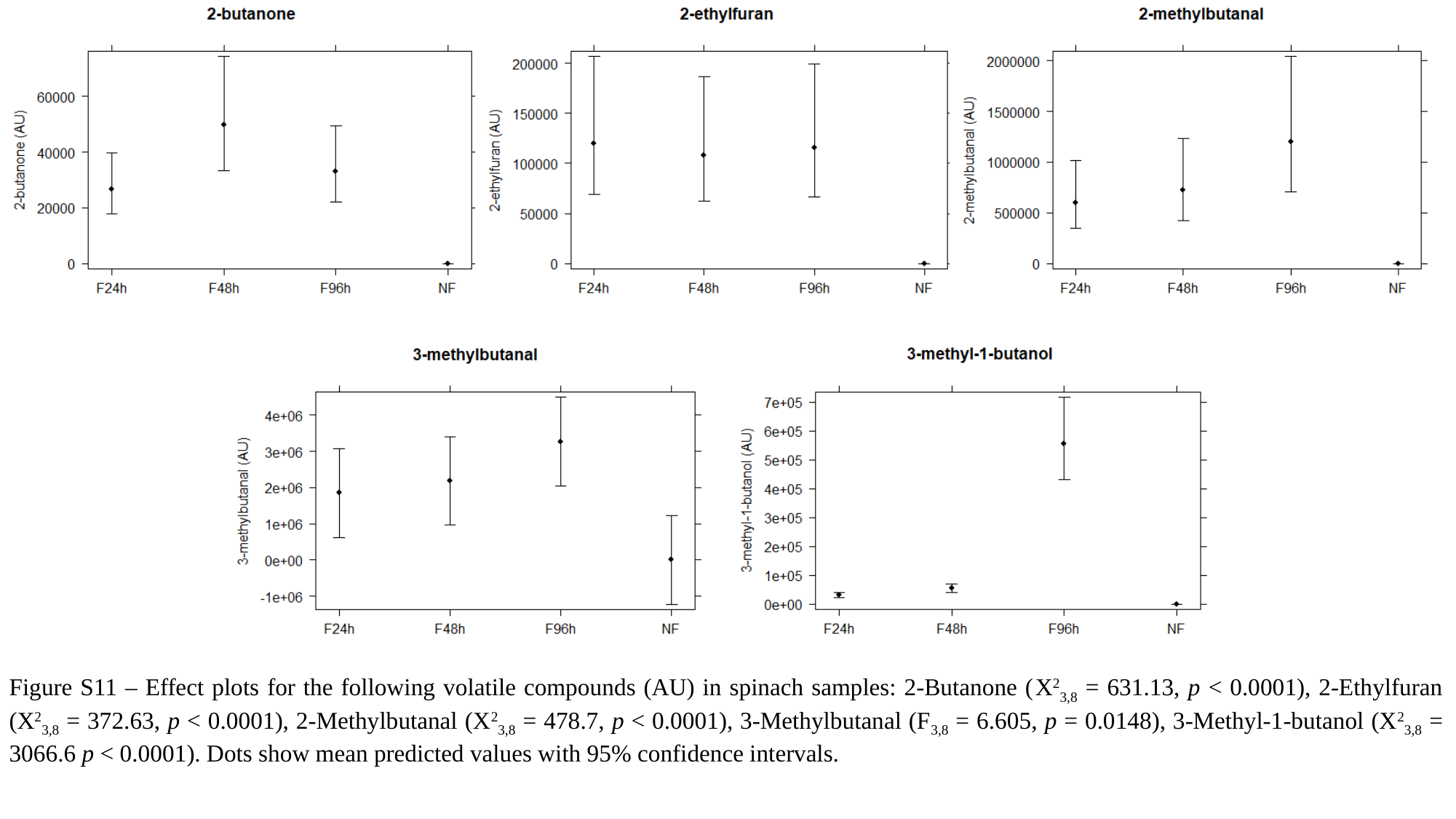

Figure S11 – Effect plots for the following volatile compounds (AU) in spinach samples: 2-Butanone (X^2^_3,8_ = 631.13, p < 0.0001), 2-Ethylfuran (X^2^_3,8_ = 372.63, p < 0.0001), 2-Methylbutanal (X^2^_3,8_ = 478.7, p < 0.0001), 3-Methylbutanal (F_3,8_ = 6.605, p = 0.0148), 3-Methyl-1-butanol (X^2^_3,8_ = 3066.6 p < 0.0001). Dots show mean predicted values with 95% confidence intervals.

### Slide 17
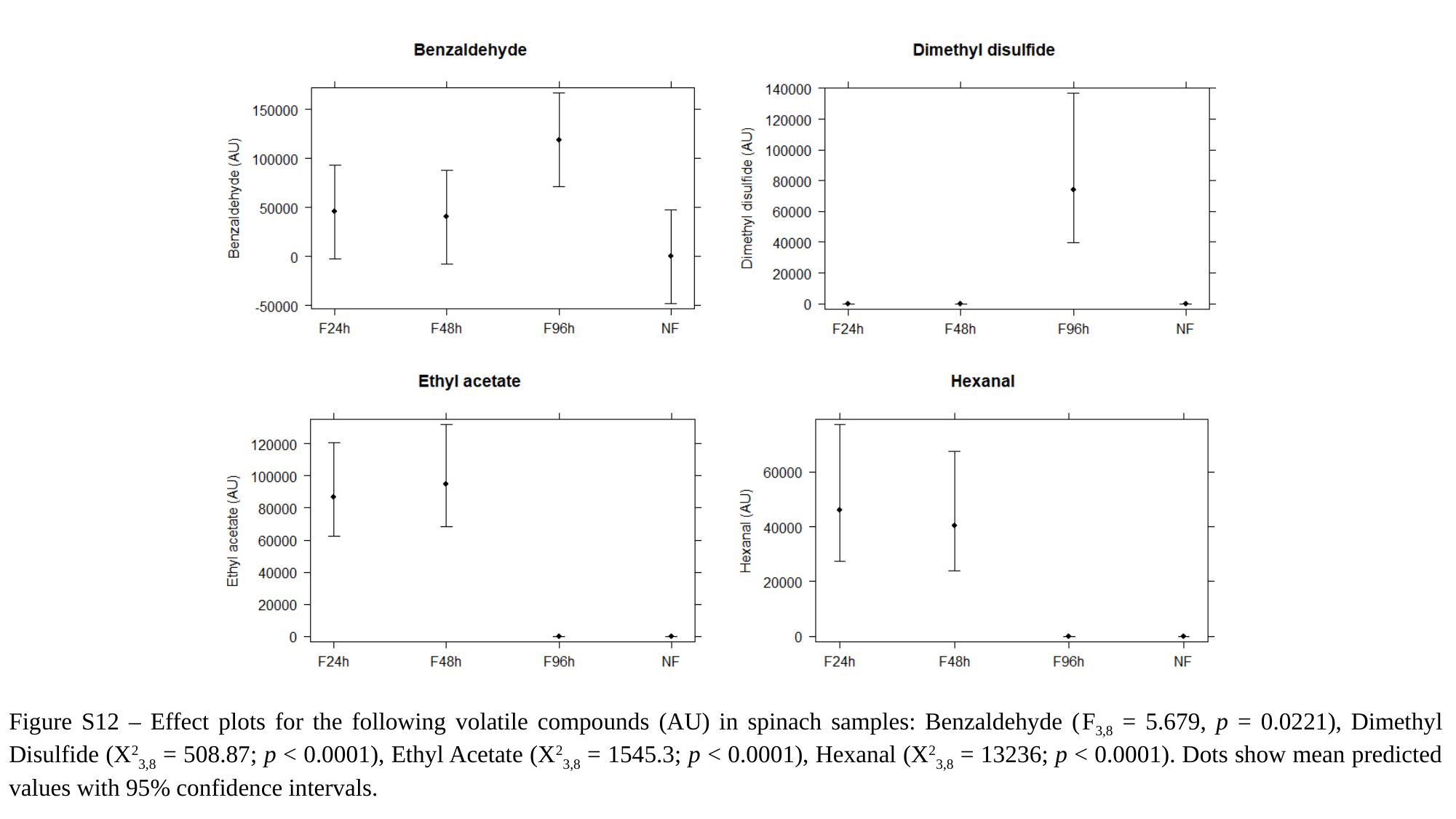

Figure S12 – Effect plots for the following volatile compounds (AU) in spinach samples: Benzaldehyde (F_3,8_ = 5.679, p = 0.0221), Dimethyl Disulfide (X^2^_3,8_ = 508.87; p < 0.0001), Ethyl Acetate (X^2^_3,8_ = 1545.3; p < 0.0001), Hexanal (X^2^_3,8_ = 13236; p < 0.0001). Dots show mean predicted values with 95% confidence intervals.

### Slide 18
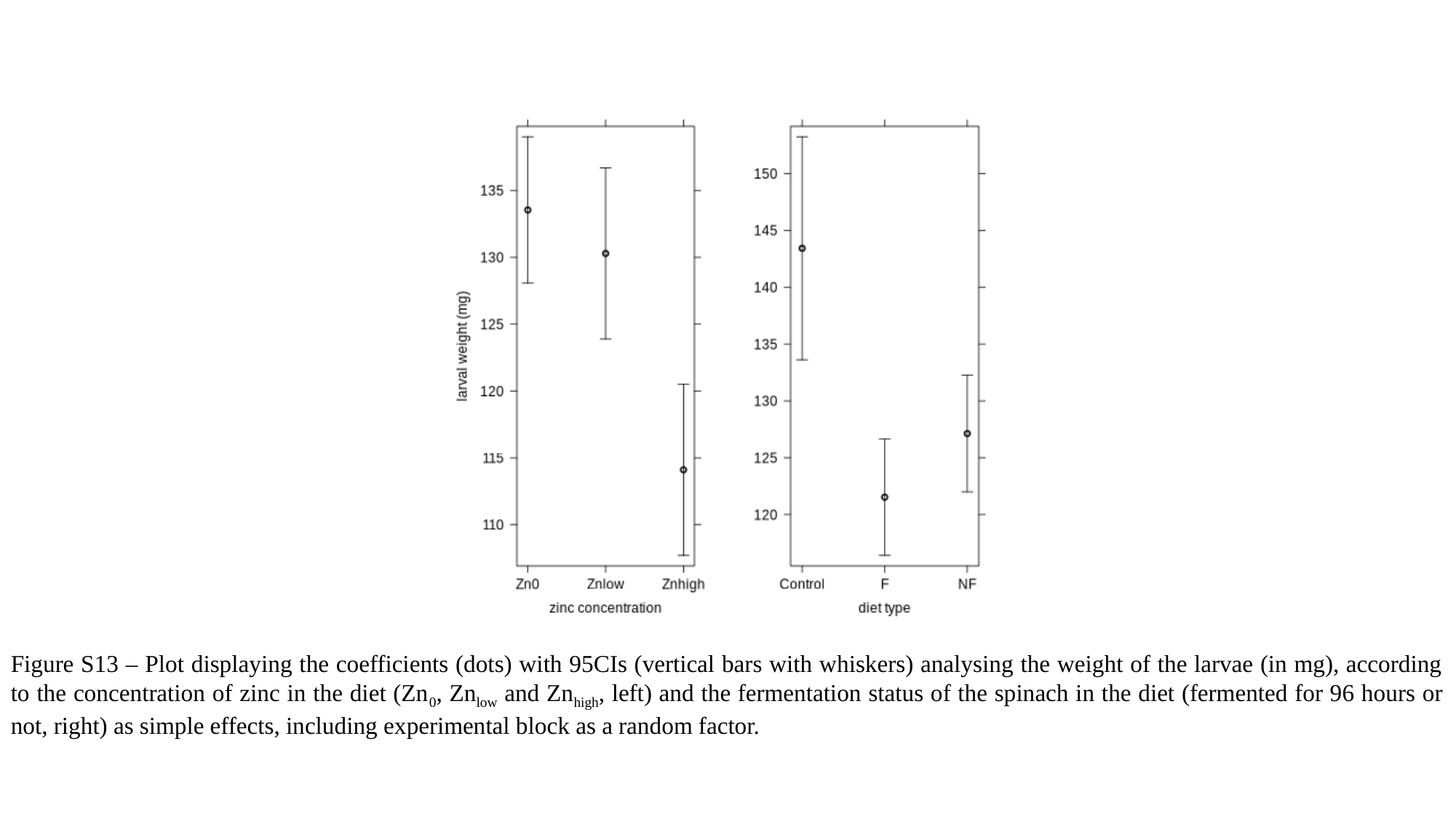

Figure S13 – Plot displaying the coefficients (dots) with 95CIs (vertical bars with whiskers) analysing the weight of the larvae (in mg), according to the concentration of zinc in the diet (Zn_0_, Zn_low_ and Zn_high_, left) and the fermentation status of the spinach in the diet (fermented for 96 hours or not, right) as simple effects, including experimental block as a random factor.

### Slide 19
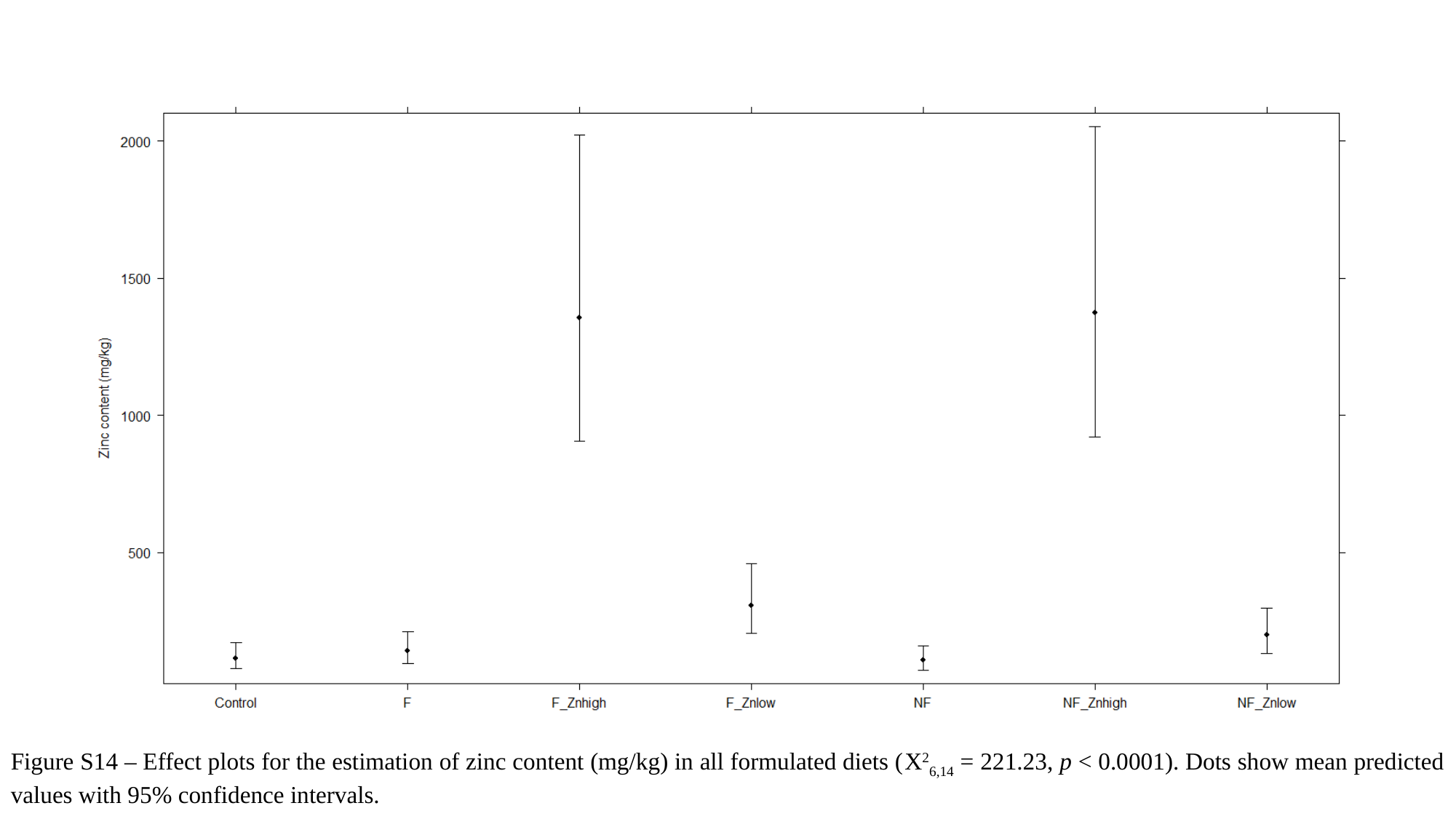

Figure S14 – Effect plots for the estimation of zinc content (mg/kg) in all formulated diets (X^2^_6,14_ = 221.23, p < 0.0001). Dots show mean predicted values with 95% confidence intervals.

### Slide 20
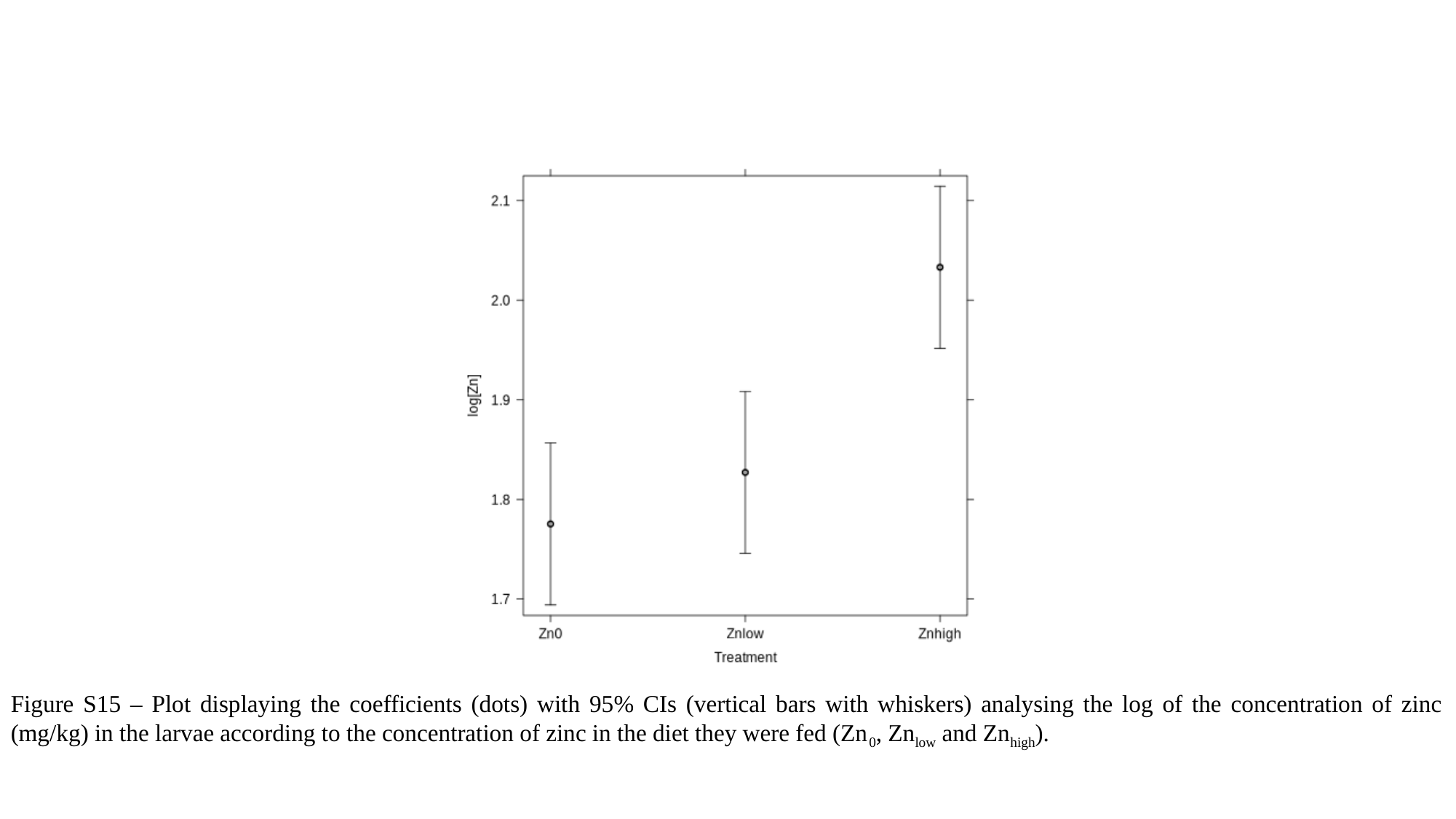

Figure S15 – Plot displaying the coefficients (dots) with 95% CIs (vertical bars with whiskers) analysing the log of the concentration of zinc (mg/kg) in the larvae according to the concentration of zinc in the diet they were fed (Zn_0_, Zn_low_ and Zn_high_).

### Slide 21
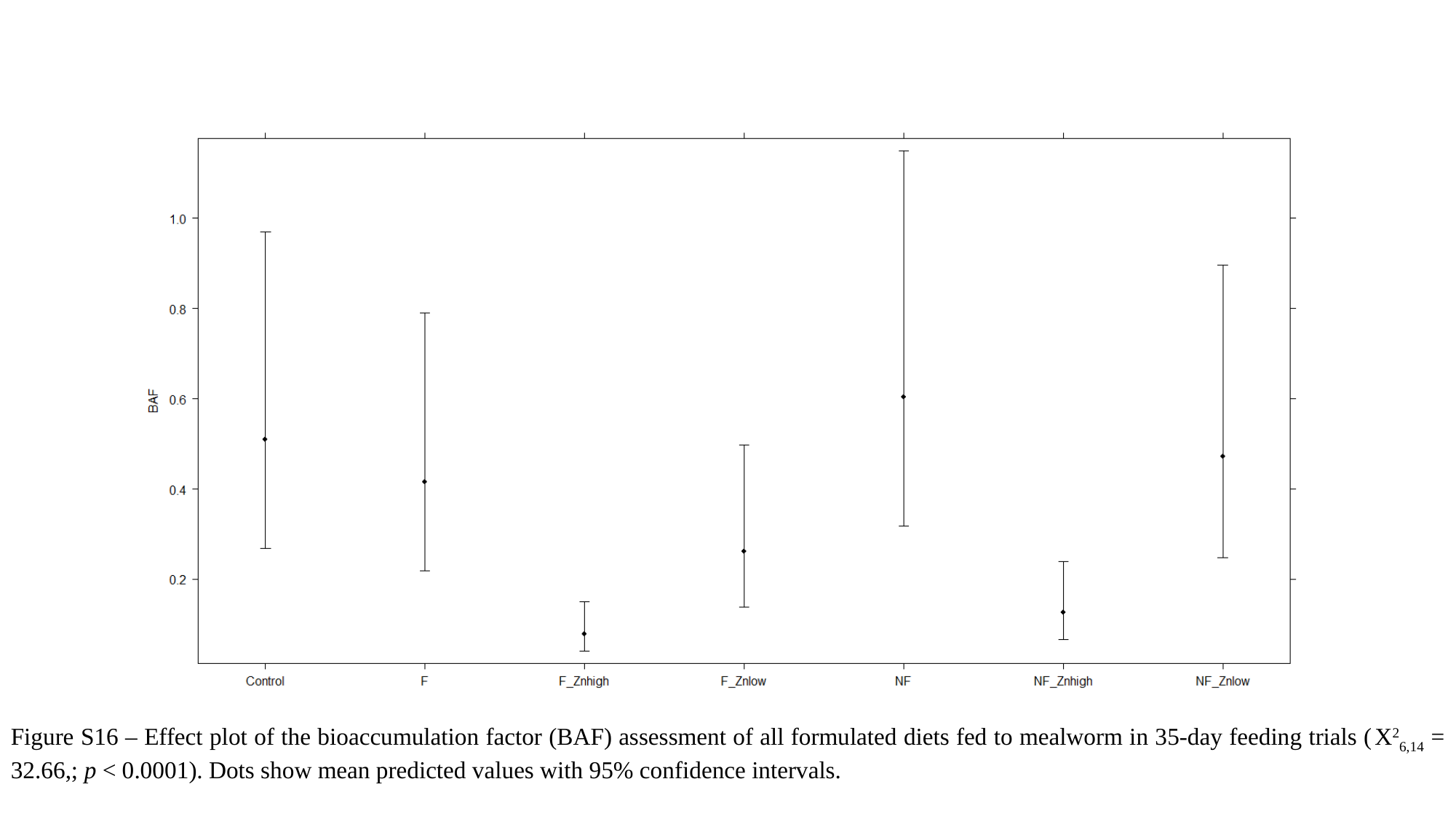

Figure S16 – Effect plot of the bioaccumulation factor (BAF) assessment of all formulated diets fed to mealworm in 35-day feeding trials (X^2^_6,14_ = 32.66,; p < 0.0001). Dots show mean predicted values with 95% confidence intervals.

### Slide 22
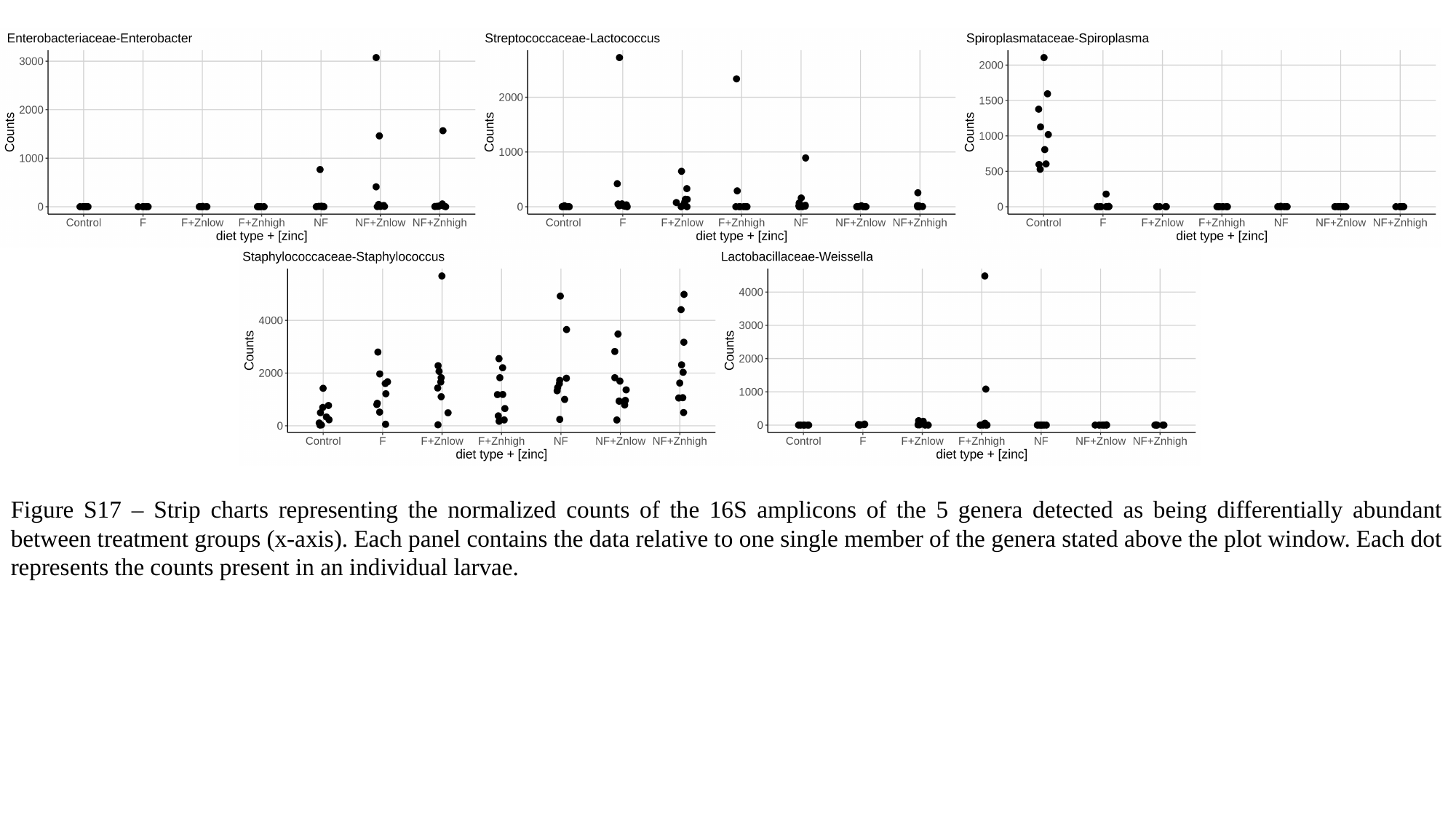

Figure S17 – Strip charts representing the normalized counts of the 16S amplicons of the 5 genera detected as being differentially abundant between treatment groups (x-axis). Each panel contains the data relative to one single member of the genera stated above the plot window. Each dot represents the counts present in an individual larvae.

### Slide 23
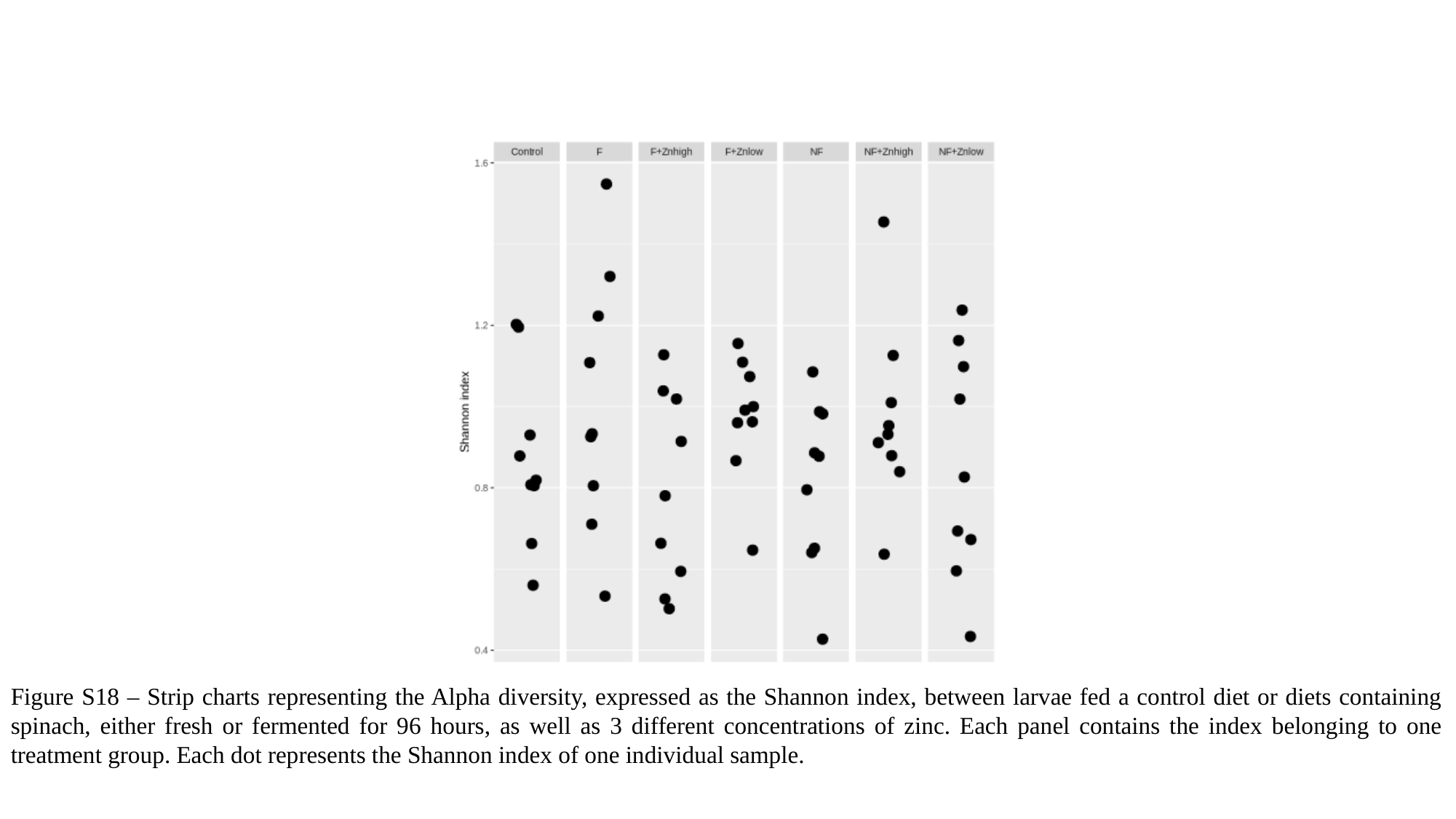

Figure S18 – Strip charts representing the Alpha diversity, expressed as the Shannon index, between larvae fed a control diet or diets containing spinach, either fresh or fermented for 96 hours, as well as 3 different concentrations of zinc. Each panel contains the index belonging to one treatment group. Each dot represents the Shannon index of one individual sample.

### Slide 24
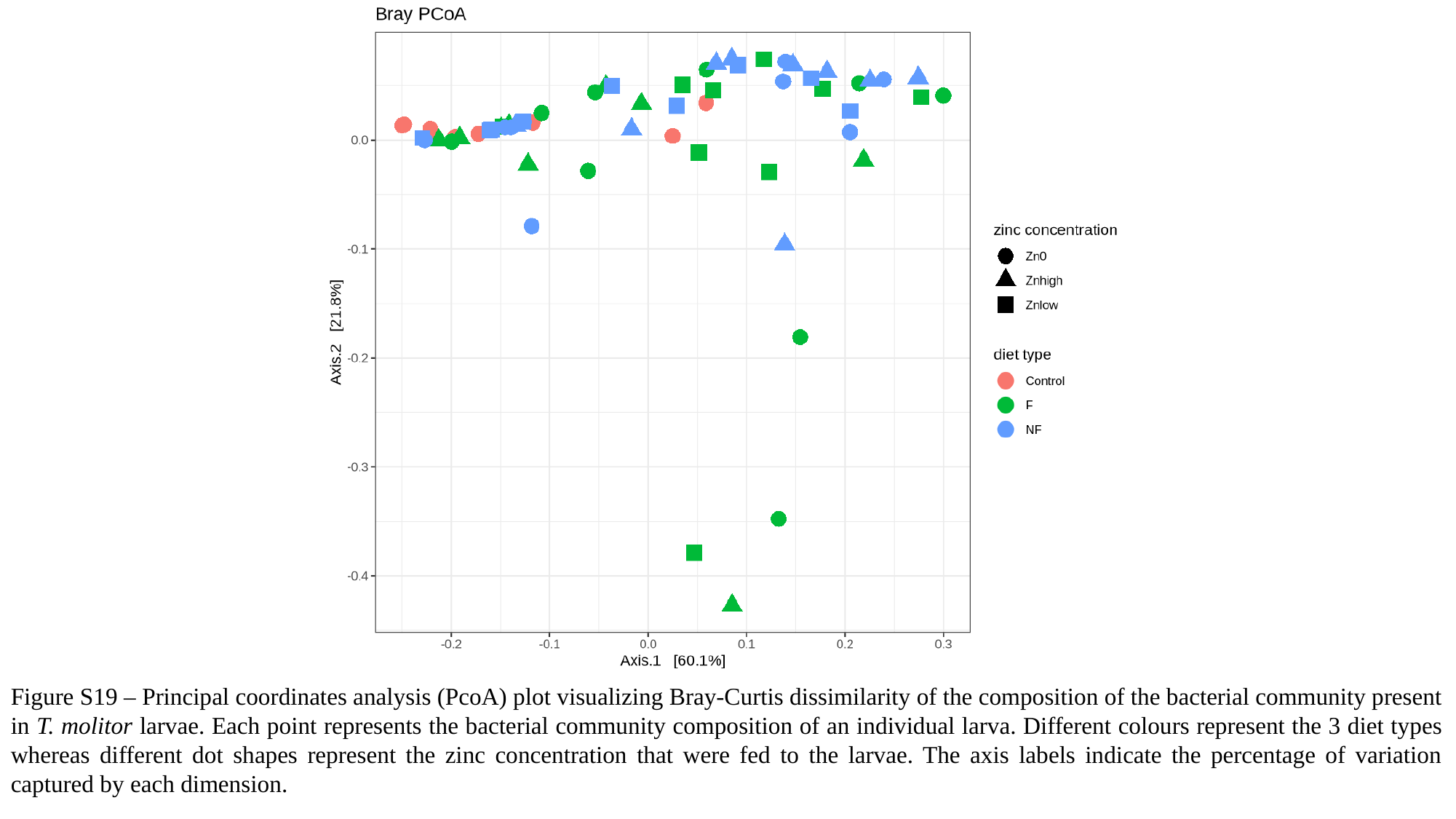

Figure S19 – Principal coordinates analysis (PcoA) plot visualizing Bray-Curtis dissimilarity of the composition of the bacterial community present in T. molitor larvae. Each point represents the bacterial community composition of an individual larva. Different colours represent the 3 diet types whereas different dot shapes represent the zinc concentration that were fed to the larvae. The axis labels indicate the percentage of variation captured by each dimension.
